# Determinants of gut microbiome composition and its response to a dietary intervention in a multi-ethnic cohort: a pop-up village study

**DOI:** 10.64898/2026.09.16.751977

**Authors:** L. Jagan, Sneha Upreti, Aditi Muglikar, Ohad Manor, Darshit Patel

**Affiliations:** Cartema Bio, Bangalore, Karnataka, India; Tovanot, NJ, USA

**Author notes:** **Corresponding Author:** Darshit Patel, Cartema Bio, No. 7/1, Haudin Rd, Halasuru, Yellappa Chetty Layout, Bangalore, Karnataka, India, 560042.

**Keywords:** Gut Microbiome, Dietary intervention, Shotgun Metagenomics, Pop-Up Village Study

## Abstract

**Background:** Dietary intervention can modulate the composition of gut microbial community. However, the consistency and magnitude of these changes may differ between individuals. Emerging evidence indicates that such variation is influenced by the baseline microbial composition and host characteristics, while variation in diet, lifestyle, and environmental exposures can further contribute to this heterogeneity. Shared-living settings, in which participants co-reside at a common site and consume a standardized diet, provide an opportunity to minimize variability arising from these external factors and to investigate the factors of microbiome composition and response to diet. In this study, we characterized the determinants of baseline gut microbial composition and evaluated the response to a short-term gut-friendly dietary intervention in a multi-ethnic cohort living under shared-living conditions.

**Methods:** Forty-three adults from North India, South India and international backgrounds co-lived in a “pop-up village” for 15 days in a single-arm design. Participants followed a gut-friendly dietary regimen. Participants were stratified by body mass index (BMI) as overweight/obese (OW, ≥ 25) or non-overweight (Non-OW, < 25). Stool samples collected at baseline and at the end of the dietary intervention (43 participants, 86 samples) were profiled by Oxford Nanopore shotgun metagenomics. Samples were evaluated for taxonomic composition, alpha diversity (per-participant linear mixed-effects models), differential abundance (consensus across three tools: MaAsLin2, LinDA, ALDEx2), and assembly-based functional gene content for six important metabolic pathways.

**Results:** At baseline, geographic region (p = 0.046) and BMI (p = 0.024) were the main factors to structure the community composition, whereas habitual diet type was not (p = 0.792). The intervention produced a small but highly reproducible shift, significant across all five distance metrics (p ≤ 0.003). The response was stronger among overweight (OW) participants, who showed a significant shift (p = 0.001), whereas non-overweight (Non-OW) participants did not (p = 0.181). The Timepoint with BMI interaction was significant for Bray-Curtis (p = 0.021) and Aitchison (p = 0.028) distances. Whole-cohort differential-abundance analysis identified depletion of the oral-associated species *Streptococcus parasanguinis* (q = 0.010) and *Gemella sanguinis* (q = 0.021), together with differential abundance of six genera, including decreased *Actinomyces* (q = 0.004) and increased *Faecalibacterium* (q = 0.016). Functional community composition of six major metabolic pathways remained largely stable at the whole-cohort level.

**Conclusion:** A short, shared gut-friendly diet elicited a modest but coordinated shift in gut microbiome composition, characterized by depletion of several oral-associated pathobiont taxa and enrichment of fibre-fermenting taxa, with the community-level compositional shift observed predominantly among overweight participants.

## 1. Introduction

The human gut microbiota is a diverse community of microorganisms primarily residing in the colon. Alterations in its composition and function have been linked with a wide range of health conditions including metabolic, inflammatory, and neurodegenerative disorders (Jackson et al., 2018; Valdes et al., 2018). The gut microbiota can be influenced by various factors such as environment, diet, geography, ethnicity, and age (Deschasaux et al., 2018; Gupta et al., 2017; Yatsunenko et al., 2012). Among these factors, diet represents one of the most influential and modifiable determinants, contributing to approximately 20% and 50% of the observed variation in microbial composition in humans and mice, respectively (Rothschild et al., 2018; Zhang et al., 2010). Therefore, understanding how dietary exposures shape microbiome composition and function remains a major focus of microbiome research.

Different dietary patterns have been shown to exert distinct effects on the composition and metabolic activity of the gut microbiota. Western-style diets characterized by high intake of fat and refined sugars have been associated with reduced microbial diversity and increased dysbiosis, whereas plant-rich and Mediterranean-style dietary patterns are associated with greater microbial diversity and beneficial metabolic profiles (Clemente-Suárez et al., 2023; De Filippis et al., 2016). In this context, increasing attention has been directed toward dietary patterns that support a beneficial gut microbial environment, often referred to as “gut-friendly” diets. These diets are typically rich in dietary fiber, whole grains, fruits, vegetables, and fermented foods, which enhance microbial diversity and stimulate the production of short-chain fatty acids, metabolites involved in maintaining intestinal homeostasis and systemic health (He et al., 2022; Holscher, 2017; Makki et al., 2018). However, dietary interventions also demonstrate substantial inter-individual variability in microbiome responses. The magnitude and direction of microbiome remodeling can depend on the pre-intervention microbial community, its functional capacity, and host metabolic characteristics (Klimenko et al., 2022; Hoffmann Sardá et al., 2025). Characterizing these inter-individual responses is challenging because dietary and environmental exposures vary substantially between individuals. Free-living observational studies, in which participants continue their normal daily lives, provide greater ecological relevance, but dietary intake and environmental exposures are difficult to control. In contrast, controlled dietary intervention studies allow researchers to tightly regulate dietary intake and, in some settings, environmental conditions, providing greater control when studying microbiome responses to specific dietary exposures (Corbin et al., 2023). However, such highly controlled settings may not fully reflect the social and environmental complexity of everyday human life.

Shared-living dietary interventions, in which participants co-reside at a single site while following a standardized dietary regimen, provide a complementary approach that combines greater control over dietary exposure with aspects of everyday living. Bringing participants together within the same setting can reduce differences in environmental exposure across individuals, potentially making dietary-associated microbiome changes easier to characterize. This approach is particularly relevant to gut microbiome research because cohabitation itself can influence microbial community structure and transmission, with cohabiting individuals showing substantial strain-level microbial sharing (Valles-Colomer et al., 2023).

In addition to these design considerations, human microbiome research remains geographically imbalanced. More than 71% of publicly available human microbiome samples with a known geographic origin come from Europe, the United States and Canada, including 46.8% from the United States alone, despite the United States representing approximately 4.3% of the global population. In contrast, India, Pakistan, and Bangladesh together account for more than one-quarter of the world’s population but contribute only 1.8% of human microbiome samples (Abdill et al., 2022). Studies conducted in India, including the LogMPIE survey of 1,004 subjects across 14 geographical locations (Dubey et al., 2018) and the multi-omics characterization of 110 healthy individuals from North-Central and Southern India by Dhakan et al. (2019), have demonstrated substantial geographic, taxonomic, and functional variation in the Indian gut microbiome. Nevertheless, controlled dietary intervention studies involving geographically and culturally diverse Indian participants remain comparatively limited.

We therefore conducted a controlled shared-living dietary intervention in a temporary co-living ‘pop-up village’ in India, bringing together adult participants from North India, South India, and international backgrounds at a single site for 15 days. Participants followed a gut-friendly diet, during which all meals were curated and provided by the study team, and external dietary intake was strictly restricted. Environmental exposures were also controlled to the extent possible to minimize external confounding factors. Within this design, we asked three key questions. First, what factors structure the gut microbiome at baseline in a multi-ethnic cohort assembled at a single site? Second, does a short, shared gut-friendly diet produce a reproducible community-level shift within a controlled shared-living setting? Third, is the magnitude of this response consistent across individuals or is it modified by host characteristics. By integrating paired taxonomic profiling, three-method consensus differential abundance, alpha-diversity modelling, and gene-content-based functional profiling, this study characterizes the microbial taxa, response direction, and host-dependent determinants associated with short-term microbiome responses following a gut-friendly dietary intervention. To our knowledge, this is the first controlled, shared-living dietary intervention conducted in a pop-up village setting in India.

## 2. Materials & Methods

### 2.1 Study design and Participant recruitment

This study was designed as a controlled, longitudinal, shared-living dietary intervention to investigate the impact of a standardized gut-friendly diet on the human gut microbiome. 43 Participants (n = 43) were recruited from diverse geographic regions, comprising from North India and South India as well as international participants, in order to capture a broad dietary and cultural diversity at baseline. All the participants lived together in a controlled “pop-up village” setting, where environmental and dietary exposures were standardized to the greatest extent possible. Participants followed a gut-friendly diet for 15 days.

Inclusion criteria included adults aged 18 years or older who were willing to comply with the dietary protocol and the sample collection schedule. Individuals with recent antibiotic use or ongoing probiotic supplementation were excluded. Participant demographic and anthropometric characteristics-age, sex, height, body weight, geographic region, country of origin, and habitual diet type (vegetarian vs non-vegetarian) were collected at each time point. A summary of these characteristics is provided in **Supplementary Table S1**. Body mass index (BMI) was calculated as body weight in kilograms divided by the square of height in metres (kg/m²) as per WHO guidelines (WHO, 1995). Participants with BMI ≥ 25 kg/m² were classified as overweight/obese (OW), and those with BMI < 25 kg/m² as non-overweight/non-obese (Non-OW).

#### 2.1.1 BMI assignment conventions

Because four participants crossed the 25 kg/m² BMI threshold between the baseline and post timepoints, we applied two complementary BMI-assignment conventions. For analyses requiring a fixed per-participant grouping across timepoints, including linear mixed-effects models, directional-consistency analysis, and core-microbiome membership, we used each participant’s baseline BMI category (baseline-BMI convention). For analyses treating each sample independently, including PERMANOVA and differential abundance, we assigned each sample to the BMI category corresponding to its own timepoint (sample-own-BMI convention). Under the sample-own convention, the pooled dataset comprised 34 OW samples and 52 Non-OW samples. At baseline, 25 participants were Non-OW and 18 OW, and at the post timepoint, 27 and 16, respectively. Geographic region (North India, South India, International) was retained as a descriptive stratification variable to examine whether baseline microbiome structure varied with geography.

### 2.2 Dietary intervention

Participants consumed a gut-friendly diet throughout the intervention period. All meals were curated and provided by the study team to ensure dietary consistency across participants. The diet primarily consisted of fiber-rich foods, whole grains, vegetables, fruits, fermented foods, and minimally processed ingredients designed to promote gut microbial health. Participants were instructed to strictly avoid outside food, alcohol, probiotics, and nutritional supplements during the intervention period. Compliance with the dietary protocol was monitored daily by the study coordinators. Individual per-participant food intake was not quantified; hence, the diet is treated as a shared dietary environment rather than an individually standardized exposure.

### 2.3 Sample collection and processing

#### 2.3.1 Sample collection

Stool samples were collected longitudinally at two predefined time points: baseline, and at the end of the intervention. Samples were self-collected by participants using Decode Biome kits, each containing an Invitek stool collection tube prefilled with DNA stabilizer (Invitek Molecular, Berlin, Germany; Cat. No. 1038111200 or 1038111300, depending on pack size), sterile sampling materials, and written instructions for proper sampling. Stool was suspended directly in the stabilizer at the point of collection. Immediately after collection, stabilized samples were transported from the study site to the laboratory at ambient temperature and stored at -80°C until DNA extraction.

#### 2.3.2 DNA extraction

Total genomic DNA was extracted from stabilized stool suspension using the QIAamp Fast DNA Stool Mini Kit (Qiagen, Hilden, Germany; cat. no. 51604) following the manufacturer’s instructions (Qiagen, 2020) with the exception that homogenized stabilized stool suspension was used as input instead of fresh stool. This stool-specific extraction protocol was selected because extraction method influences recovered community composition and downstream association analyses (Fernández-Pato et al., 2024).

Extracted DNA was subjected to a bead-based clean-up with AMPure XP magnetic beads (Beckman Coulter, Brea, CA, USA; cat. no. A63881) to remove short DNA fragments, residual salts, and enzymatic carry-over (Beckman Coulter, 2016). Bound DNA was washed twice with freshly prepared 80 percent ethanol and eluted in the Acetate-Tris-EDTA (ATE) buffer supplied with the extraction kit.

DNA concentration and purity were assessed prior to library preparation. Purity was evaluated on a NanoDrop spectrophotometer (Thermo Fisher Scientific, Waltham, MA, USA) using the A260/A280 absorbance ratio, with samples accepted at a ratio of 1.8-2.0. DNA concentration was quantified fluorometrically using the Qubit dsDNA HS Assay Kit on a Qubit fluorometer (Thermo Fisher Scientific; Thermo Fisher Scientific, 2015).

#### 2.3.3 Library preparation and sequencing

Sequencing libraries were prepared from the purified genomic DNA using the Native Barcoding Kit SQK-NBD114 (Oxford Nanopore Technologies plc, Oxford, UK) according to the manufacturer’s protocol (Oxford Nanopore Technologies, 2023). Barcoded libraries were pooled and loaded onto PromethION R10.4.1 flow cells. Sequencing was performed on a PromethION 2 Solo (P2 Solo) device under the control of MinKNOW, and raw signal data were basecalled with Dorado using the high-accuracy (hac) model (Dorado7.6.7) (MinKNOW 24.11.8). Long-read shotgun metagenomic sequencing was selected for its capacity to resolve microbial taxa at high taxonomic resolution from complex fecal communities (Marić et al., 2024).

Across all sequencing runs, the workflow generated approximately 153.2 Gb of sequence data. Per-sample mean read length ranged from 1.4 to 5.5 kb, and per-sample data output ranged from 512 Mb to 5.05 Gb.

### 2.4 Bioinformatics analysis

Raw sequencing reads were trimmed of adapter sequences and split at internal adapters (chimera detection) with Porechop v0.2.4 (rrwick/Porechop; default parameters). Reads shorter than 800 bp and with Q-score below 7 were discarded. Host-decontaminated reads were classified taxonomically with Kraken2 against the HumGut database (gutseq v2.0, released in July 2025). Kraken2 read counts assigned at species and genus level were retained as abundance estimates.

For functional profiling, decontaminated reads were assembled with Flye in --meta mode, and short contigs unlikely to represent real genomes were removed. Ribosomal and transfer RNA regions were masked with Barrnap and Aragorn, respectively, to prevent Prodigal from calling spurious ORFs inside rRNA and tRNA loci. Protein-coding genes were predicted with Prodigal on the masked assembly in metagenomic mode and searched with DIAMOND blastp (e-value 0.001) against the KEGG GENES database (release Sep 3, 2023), with orthology assigned by KOfam profile HMMs. Reads were mapped back to the filtered contigs with minimap2 and samtools, and per-position coverage was combined with gene coordinates and contig lengths to quantify each KEGG function per sample and aggregate them into pathway-level summaries.

### 2.5 Statistical analysis

All analyses were performed in R (4.6.1) using phyloseq. The primary phyloseq object comprised 1,131 taxa across 86 samples resolved at seven taxonomic ranks, with an accompanying phylogenetic tree.

#### 2.5.1 Data processing and normalization

Count data were total-sum-scaled to relative abundances and centered-log-ratio (CLR) transformed for compositional analyses.

#### 2.5.2 Diversity analyses

Six alpha-diversity indices were computed per sample: observed richness, Chao1, Shannon and Simpson indices, Pielou’s evenness (all via the microbiome package), and Faith’s phylogenetic diversity (picante). Timepoint effects were tested with per-metric linear mixed-effects models fitted with nlme::lme (fixed effect: timepoint; random effect: participant intercept); marginal and conditional R² were computed following Nakagawa & Schielzeth as implemented in MuMIn (Bartoń, 2023). Analyses were performed separately in the whole cohort and stratified by baseline BMI category; p-values were adjusted for the six metrics within each analysis using the Benjamini–Hochberg procedure. Beta diversity was computed using Bray-Curtis, Jaccard, weighted and unweighted UniFrac, and Aitchison distances and visualized with principal-coordinates analysis (PCoA), including paired PCoA plots linking each participant’s timepoints. PERMANOVA was performed with vegan::adonis2 using 999 permutations with permutations constrained within participant (strata=Participant). Timepoint × BMI, Timepoint × Region, and Timepoint × Diet interactions were each tested in a separate model containing Timepoint, the respective covariate main effect, and their interaction. Baseline determinants (region, BMI, and diet type) were tested by cross-sectional PERMANOVA on the 43 baseline samples without strata. Homogeneity of multivariate dispersion was assessed with betadisper, and taxa contributing to between-group dissimilarity were identified with SIMPER. Core-microbiome membership was defined by prevalence and reported separately for each BMI stratum, using a threshold of ≥ 0.1% relative abundance in ≥ 50% of participants per timepoint.

#### 2.5.3 Association testing

The same statistical framework was applied to taxonomic features (species and genus) and to functional features (KOs and their parent pathways). Paired timepoint comparisons used the Wilcoxon signed-rank test with Benjamini-Hochberg correction. Per-feature linear mixed-effects models were fitted with nlme::lme using a Participant random intercept to test Timepoint, BMI, and their interaction; marginal and conditional R² computed following Nakagawa & Schielzeth as implemented in MuMIn, and interaction terms were evaluated by likelihood-ratio tests. Differential abundance of taxa was evaluated using a three-tool consensus spanning complementary modelling assumptions: MaAsLin2 (with Participant as a random effect), LinDA (a mixed-model extension for compositional data), and ALDEx2 (Monte Carlo Dirichlet sampling with CLR transformation); features were assigned to confidence tiers according to the number of tools in agreement. KO-level differential abundance was tested with MaAsLin2 under the same paired design specification. Interaction effects were further interrogated with per-participant binomial directional-consistency tests and with label-shuffling permutation.

## 3. Results

A consolidated summary is provided in **Figure 1**, with the underlying analyses reported in the subsections that follow.

**Fig. 1.**
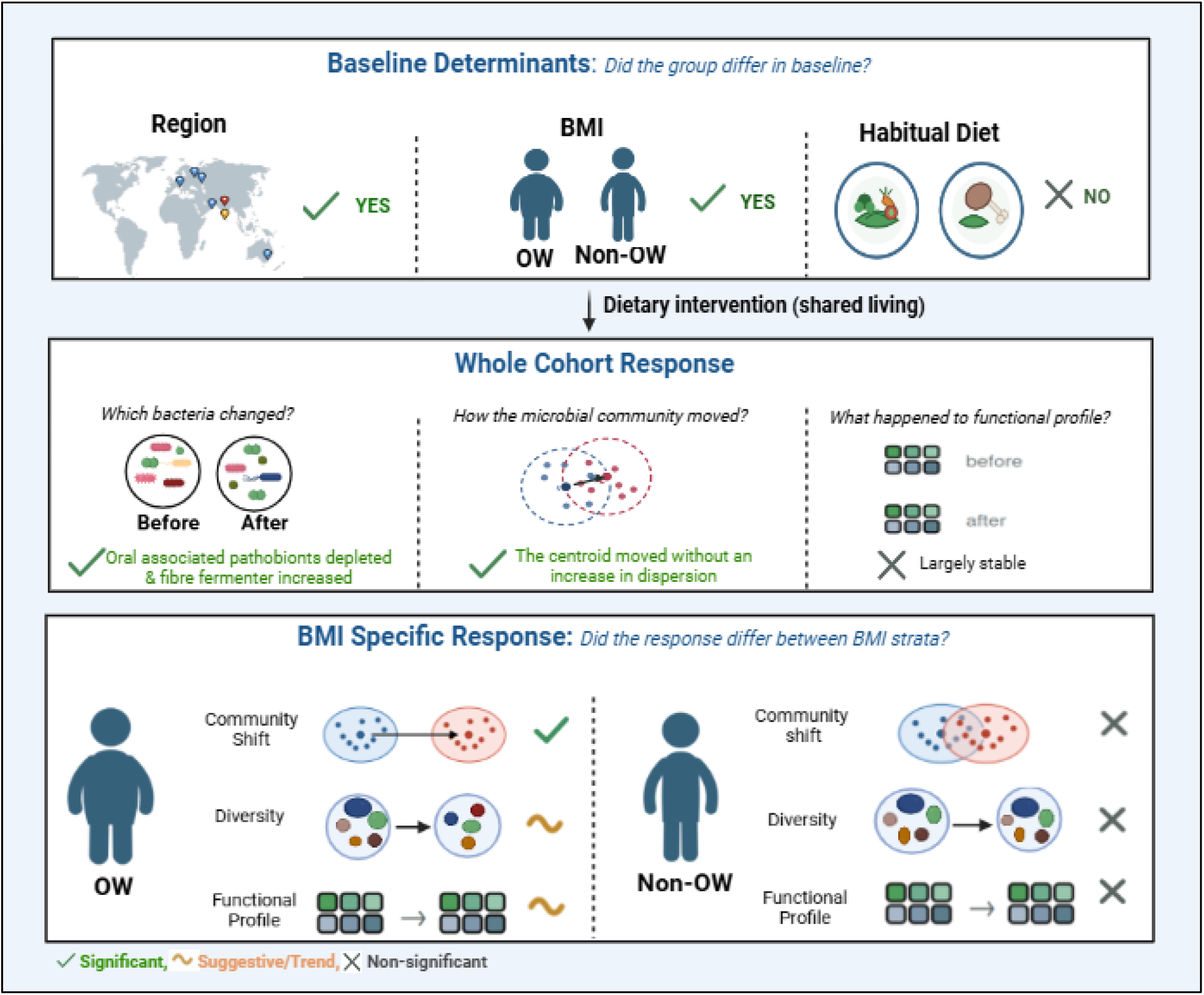
Consolidated summary of the main findings. Top panel. (Baseline Determinants) shows region and BMI were associated with baseline composition and diet type was not. Pin color shows different region (red = North India, yellow = South India, blue = International). **Middle panel** (Whole-cohort response) shows the effect of intervention at whole cohort level. **Bottom panel** (BMI-specific response) shows the effect of intervention in BMI specific strata; only overweight (OW) participants showed a community-level shift, with trend-level changes in diversity and function. Tick = significant; tilde = trend-level; cross = not significant. **Created with** BioRender.com.

### 3.1 Geographic region and BMI, not the diet type, structure the baseline composition of the gut microbiome

We first characterized the factors that shape the gut microbiome at baseline, before any dietary intervention. Cross-sectional PERMANOVA of 43 participants (43 baseline samples) showed that microbial taxonomic composition was significantly associated with geographic region and BMI, whereas diet type (habitual dietary pattern, recorded as vegetarian or non-vegetarian) showed no significant effect. Geographic region accounted for the largest proportion of variation in microbial composition (Bray-Curtis, R² = 6.42%, *p* = 0.046), followed by BMI (R² = 4.00%, *p* = 0.024). These associations were consistently observed across weighted UniFrac and Aitchison distance metrics, indicating that the findings were robust to different measures of microbial community dissimilarity.

In contrast, diet type explained only a small and non-significant proportion of the variance under all three metrics (Bray-Curtis R² = 1.88%, *p* = 0.792; Weighted UniFrac R² = 1.93%, p = 0.796; Aitchison R² = 2.40%, p = 0.398). Beta dispersion tests for region (*p* = 0.400) and BMI (*p* = 0.692) were non-significant, indicating that the significant PERMANOVA results for these factors reflected true differences in community composition rather than differences in within-group variability **(Table 1**, **Figure 2)**.

**Table 1.** Baseline PERMANOVA (n = 43) for region, BMI, and diet. Each row is a separate PERMANOVA model testing one host factor under one distance metric. R² = proportion of total variation explained by the factor; P = permutation p-value; Betadisper p = homogeneity-of-dispersion p-value.

| Factor | Metric | $R^2$ | P | Betadisper p |
| --- | --- | --- | --- | --- |
| Region | Bray-Curtis | 6.42% | 0.046 | 0.400 |
| Region | Weighted UniFrac | 6.29% | 0.044 | - |
| Region | Aitchison | 6.84% | 0.017 | - |
| BMI | Bray-Curtis | 4.00% | 0.024 | 0.692 |
| BMI | Weighted UniFrac | 4.00% | 0.015 | - |
| BMI | Aitchison | 3.74% | 0.021 | - |
| Diet | Bray-Curtis | 1.88% | 0.792 | 0.906 |
| Diet | Weighted UniFrac | 1.93% | 0.796 | - |
| Diet | Aitchison | 2.40% | 0.398 | - |

**Fig. 2.**
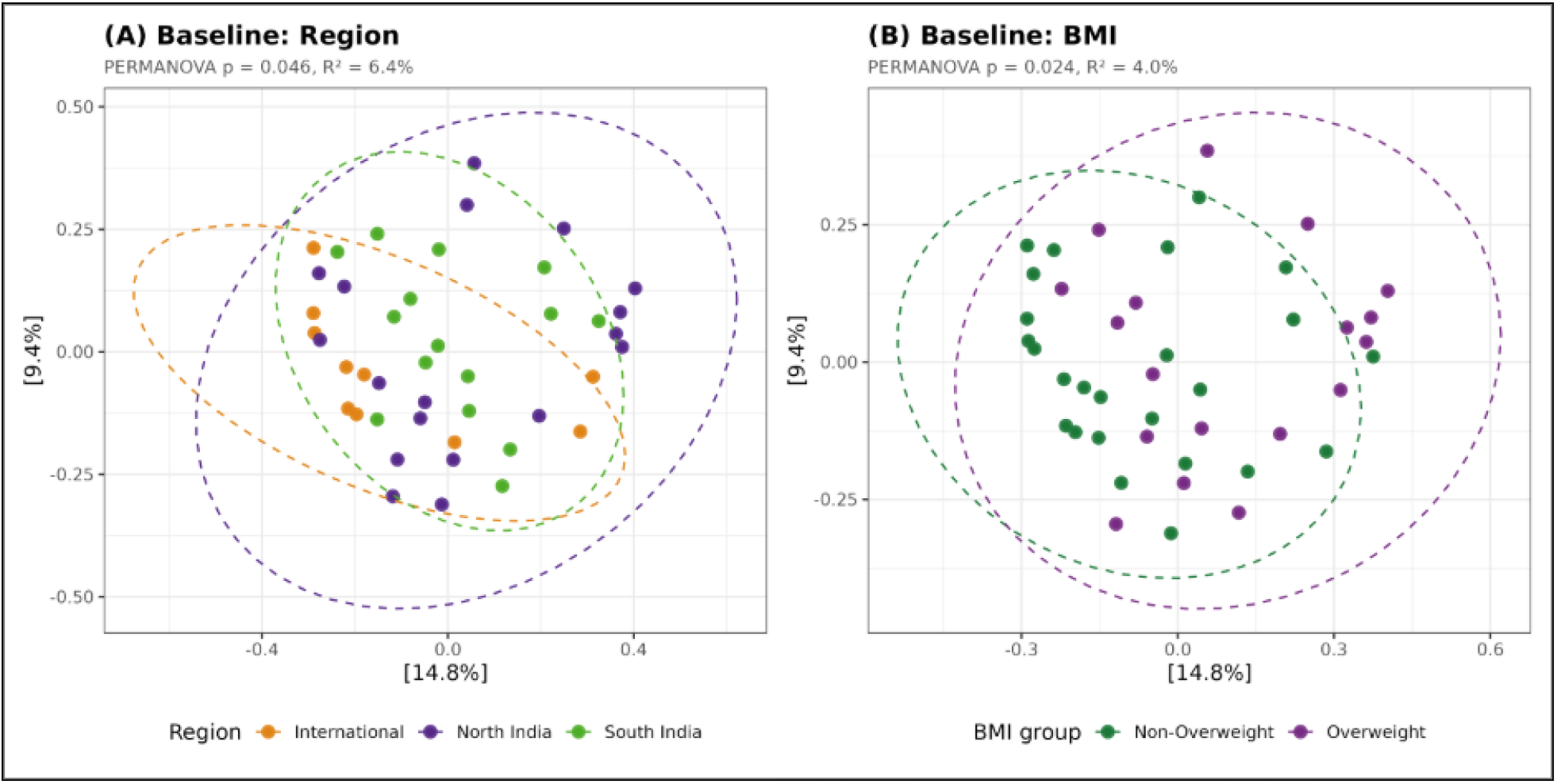
Baseline gut microbial community structure by geography and BMI. Principal coordinates analysis (PCoA) of baseline fecal metagenomes (Bray-Curtis dissimilarity, TSS-normalized). **(A)** Samples colored by region (International n=10, North India n=18, South India n=15); **(B)** Samples colored by BMI group (Non-OW n=25, OW n=18). Each point is one participant at baseline; ellipses denote 95% confidence regions per group.

### 3.2 A short term dietary intervention produces a coordinated shift at the cohort level

We further tested whether the shared gut-friendly diet produced a detectable community-level shift. Paired PERMANOVA was applied to each participant’s baseline and post-intervention samples under five distance metrics. All five yielded a significant timepoint effect (Aitchison R² = 1.15%, Bray-Curtis R^2^ = 1.40%, weighted UniFrac R² =1.41%, Jaccard R² = 1.46%, unweighted UniFrac R² = 1.58%; all p ≤ 0.003; **Table 2**, **Figure 3**). Although the effect size was small (R² = 1.15-1.58%), the consistent timepoint effect across all five distance metrics supports a reproducible shift in community composition following the intervention.

**Table 2.**
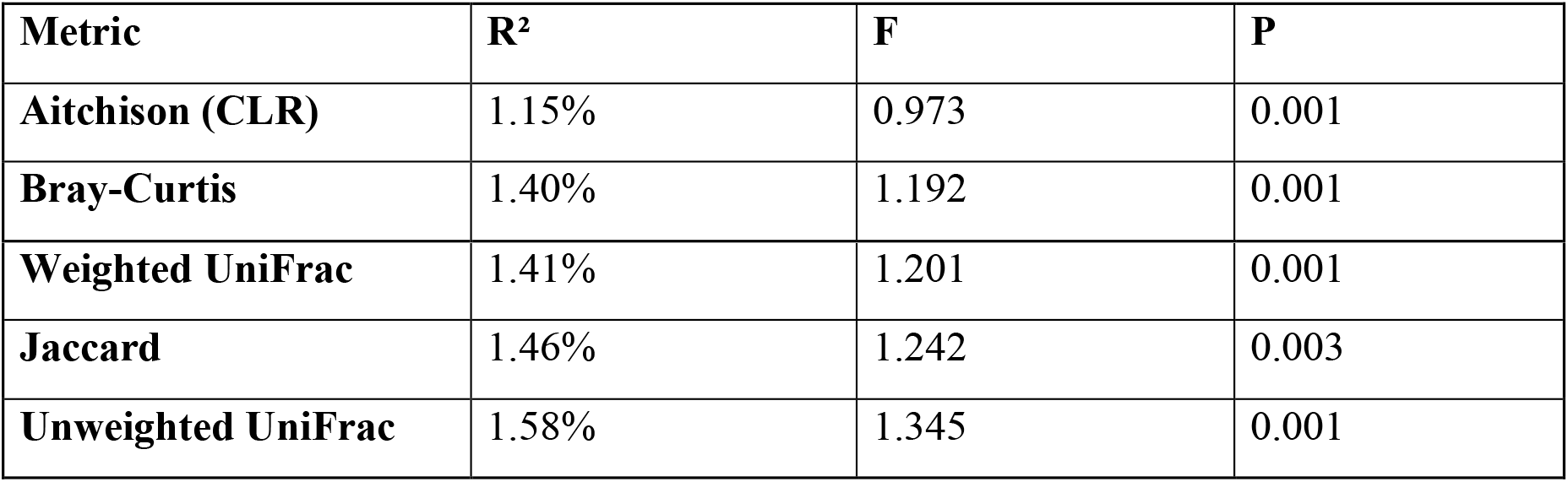
Paired PERMANOVA: Timepoint effect across five distance metrics (n=43, 86 samples). Each row is a separate PERMANOVA model testing the difference in community composition between baseline and post-intervention samples. R² = proportion of total variation explained by timepoint; F = pseudo-F ratio; P = permutation p-value.

| Metric | $R^2$ | F | P |
| --- | --- | --- | --- |
| Aitchison (CLR) | 1.15% | 0.973 | 0.001 |
| Bray-Curtis | 1.40% | 1.192 | 0.001 |
| **Weighted UniFrac** | 1.41% | 1.201 | 0.001 |
| **Jaccard** | 1.46% | 1.242 | 0.003 |
| **Unweighted UniFrac** | 1.58% | 1.345 | 0.001 |

**Fig. 3.**
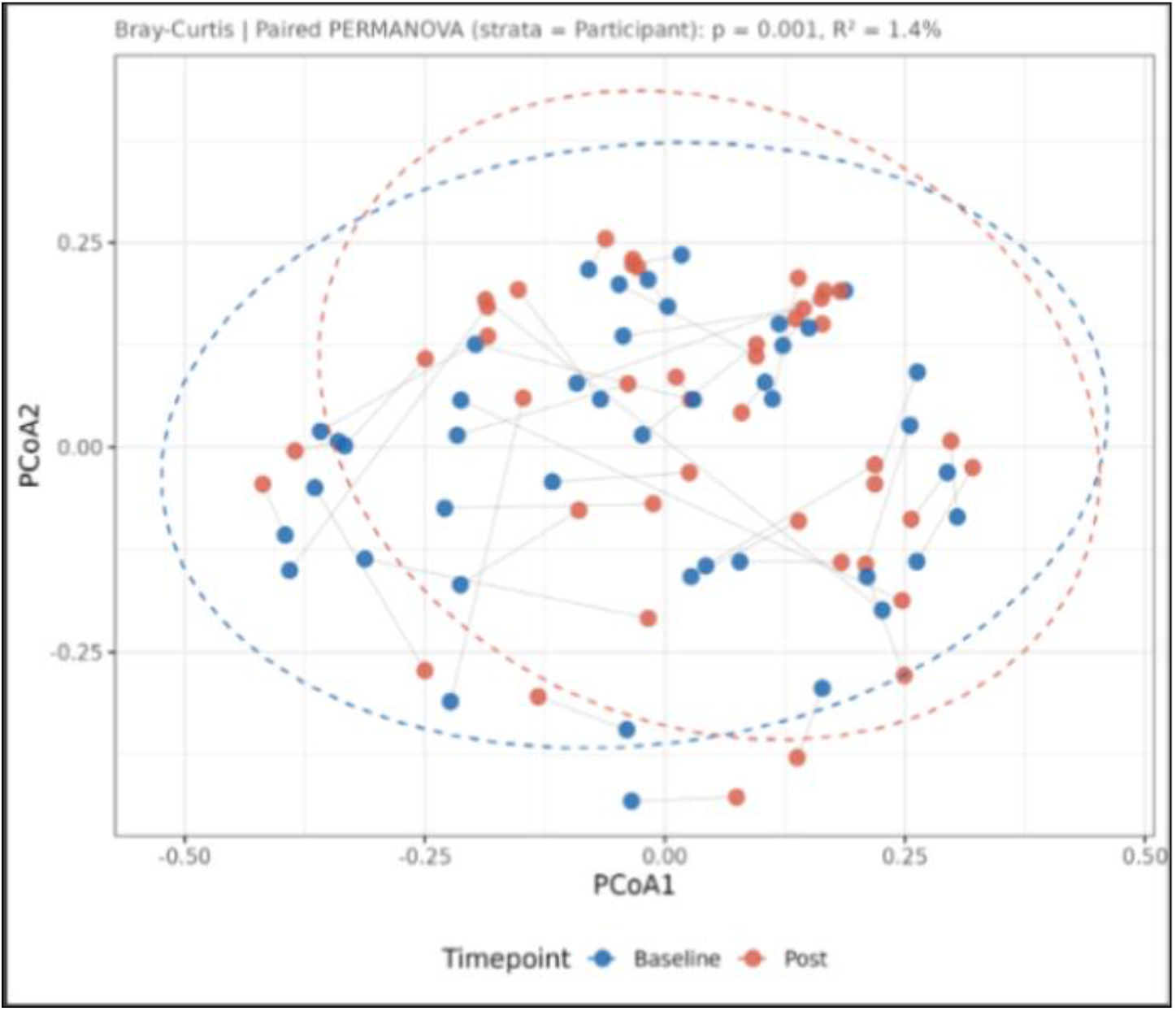
Short-term dietary intervention shifts overall gut microbial community composition. Principal coordinates analysis (PCoA) of paired baseline (blue) and post-intervention (red) fecal metagenomes (86 samples, 43 participants; grey lines connect each participant’s baseline and post-intervention samples in the ordination space. Bray-Curtis dissimilarity, TSS-normalized). Ellipses denote 95% confidence regions per timepoint. Paired PERMANOVA (strata = Participant, 999 permutations): p = 0.001, R² = 1.4%.

A significant PERMANOVA effect indicates a difference in overall community composition between timepoints, but does not, by itself, distinguish whether this difference reflects a shift in community centroid or increased inter-individual dispersion. A centroid shift represents a change in the overall position of the community while maintaining relatively similar dispersion among samples, whereas increased dispersion reflects greater variability among participants without a substantial shift in the overall centroid (**Figure 4**).

**Fig. 4.**
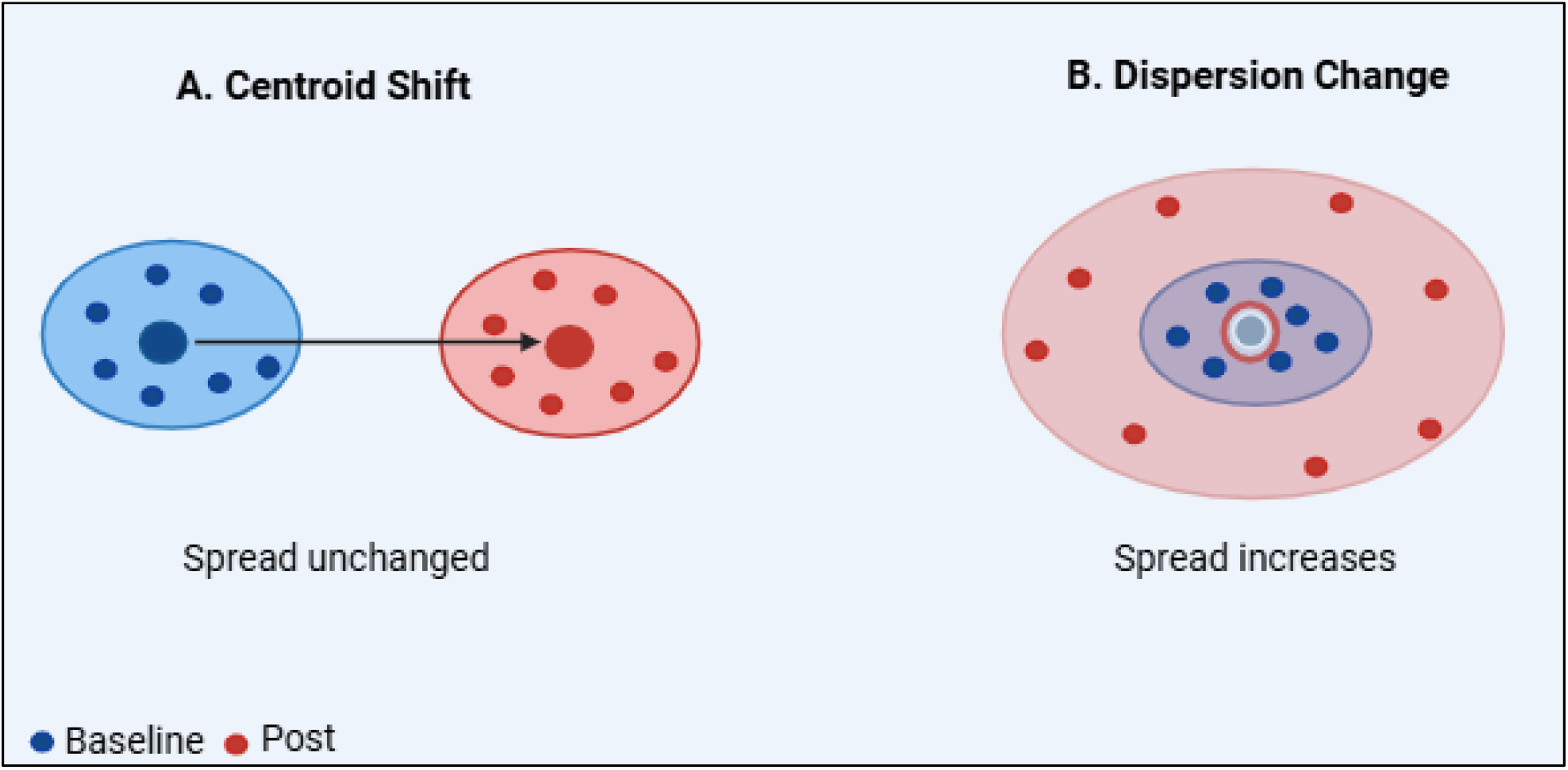
Schematic illustration of two possible shifts underlying a significant PERMANOVA effect. Illustrative images; blue = baseline (B), red = post-intervention (P); **(A)** Centroid shift: the centroid moves while within-group spread is unchanged. **(B)** Dispersion change: the centroid stays in place while within-group spread increases. **Created with BioRender.com.**

To distinguish between these possibilities, we assessed homogeneity of multivariate dispersion using betadisper. Dispersion did not differ significantly between baseline and post-intervention samples across any of the five distance metrics (p = 0.182 to 0.806; **Supplementary Table S2**). A complementary within-participant comparison of distances to the group centroid was also non-significant (Wilcoxon signed-rank test, V = 561, p = 0.291). Together, these findings provide no evidence for increased inter-individual dispersion and are consistent with the significant PERMANOVA effect reflecting a shift in community centroid rather than increased heterogeneity among participants, the scenario depicted in **Figure 4A**.

### 3.3 BMI modifies the response to dietary intervention

We further examined whether the overall shift was uniform across the cohort or driven by a specific subgroup. Three separate paired PERMANOVA models were fitted on Bray-Curtis distances, testing the Timepoint × BMI, Timepoint × Region and Timepoint × Diet interactions independently. Only the Timepoint × BMI interaction was significant (R² = 1.16%, p = 0.021), the Timepoint × Region (R² = 0.76%, p = 0.893) and Timepoint × Diet (R² = 0.50%, p = 0.483) interactions were not **(Table 3).** This interaction was reproducible under the compositional Aitchison distance (p = 0.028) and showed a concordant, non-significant trend under weighted UniFrac (p = 0.095; **Supplementary Table S3**). This indicates that host BMI, rather than geography or diet, modified the microbiome response to the intervention.

**Table 3.** Bray-Curtis paired PERMANOVA (86 samples). Each model included Timepoint, the corresponding covariate main effect (BMI, Region, or Diet), and their interaction. Main effects are not reported as this analysis addresses the interaction terms; residual df and R² reflect their inclusion.

| Term | Df | SumOfSqs | R<sup>2</sup> | F | P |
| --- | --- | --- | --- | --- | --- |
| **Model 1** |  |  |  |  |  |
| Timepoint | 1 | 0.357 | 1.40% | 1.204 | 0.001 |
| Timepoint × BMI | 1 | 0.296 | 1.16% | 0.998 | 0.021 |
| Residual | 82 | 24.309 | 95.29% |  |  |
| **Model 2** |  |  |  |  |  |
| Timepoint | 1 | 0.357 | 1.40% | 1.205 | 0.001 |
| Timepoint × Region | 2 | 0.195 | 0.76% | 0.329 | 0.893 |
| Residual | 80 | 23.689 | 92.86% |  |  |
| **Model 3** |  |  |  |  |  |
| Timepoint | 1 | 0.357 | 1.40% | 1.193 | 0.001 |
| Timepoint × Diet | 1 | 0.128 | 0.50% | 0.429 | 0.483 |
| Residual | 82 | 24.523 | 96.13% |  |  |

Habitual diet type was associated with neither baseline community structure (Section 3.1, p = 0.792) nor the response to the intervention (Timepoint × Diet, p = 0.483). Geographic region structured the community at baseline but likewise did not modify the response. BMI was the only host characteristic that both structured the community at baseline and modified the response to the intervention (**Table 4**).

**Table 4.** Host factors tested against baseline community composition and against the response to the intervention. Baseline column represents cross-sectional PERMANOVA on the 43 baseline samples. The response column represents Timepoint × factor interaction from paired PERMANOVA on all 86 samples.

| Host Factor | Baseline composition | Response to intervention |
| --- | --- | --- |
| Habitual diet type | Non-significant ($p = 0.792$) | Non-significant ($p = 0.483$) |
| Geographic region | Significant ($p = 0.046$) | Non-significant ($p = 0.893$) |
| BMI | Significant ($p = 0.024$) | Significant ($p = 0.021$) |

BMI-stratified analysis confirmed that the shift was restricted to OW participants (Bray-Curtis R² = 4.92%, p = 0.001; weighted UniFrac R² = 4.63%, p = 0.001) and was non-significant in Non-OW participants (Bray-Curtis p = 0.181; **Table 5, Supplementary Figure S1).** The taxa contributing specifically to this OW shift are ranked in **Supplementary Table S4** and **Supplementary Figure S2.**

**Table 5.** BMI-stratified paired PERMANOVA. BMI assignments by each sample’s own BMI at collection. A significant timepoint effect was present among OW participants but not among Non-OW participants.

| Subgroup | Metric | R<sup>2</sup> | F | P |
| --- | --- | --- | --- | --- |
| **OW (34 samples)** | Bray-Curtis | 4.92% | 1.657 | 0.001 |
| **OW (34 samples)** | Weighted UniFrac | 4.63% | 1.552 | 0.001 |
| **Non-OW (52 samples)** | Bray-Curtis | 1.10% | 0.555 | 0.181 |
| **Non-OW (52 samples)** | Weighted UniFrac | 1.17% | 0.590 | 0.177 |

To determine whether the more pronounced microbiome shift in participants with OW reflected larger individual changes or a more consistent direction of change across individuals, we compared within-subject community changes between the BMI groups. The magnitude of change, quantified as the per-participant baseline-to-post-intervention, Bray-Curtis dissimilarity extracted from the full distance matrix, did not differ significantly between the OW (mean = 0.527; n = 18, classified by baseline BMI) and Non-OW groups (mean = 0.460; n = 25; Wilcoxon rank-sum test, p = 0.196, r = 0.236; **Figure 5A**). Consistent with this, the lengths of individual PCoA trajectories were comparable between groups (arrow-length permutation test, p = 0.273; **Figure 5B**), and within-subject dissimilarities showed no significant variation across geographic regions (International 0.448; North India 0.492; South India 0.510; Kruskal-Wallis test, p = 0.675) or dietary groups (p = 0.667). In contrast, the displacement of the group centroid from baseline to post-intervention was substantially greater in the OW group (0.154) than in the Non-OW group (0.012; **Figure 5C**). Together, these results indicate that the more pronounced shift associated with OW arose from greater consistency in the direction of individual community changes rather than larger within-subject compositional changes.

**Fig. 5.**
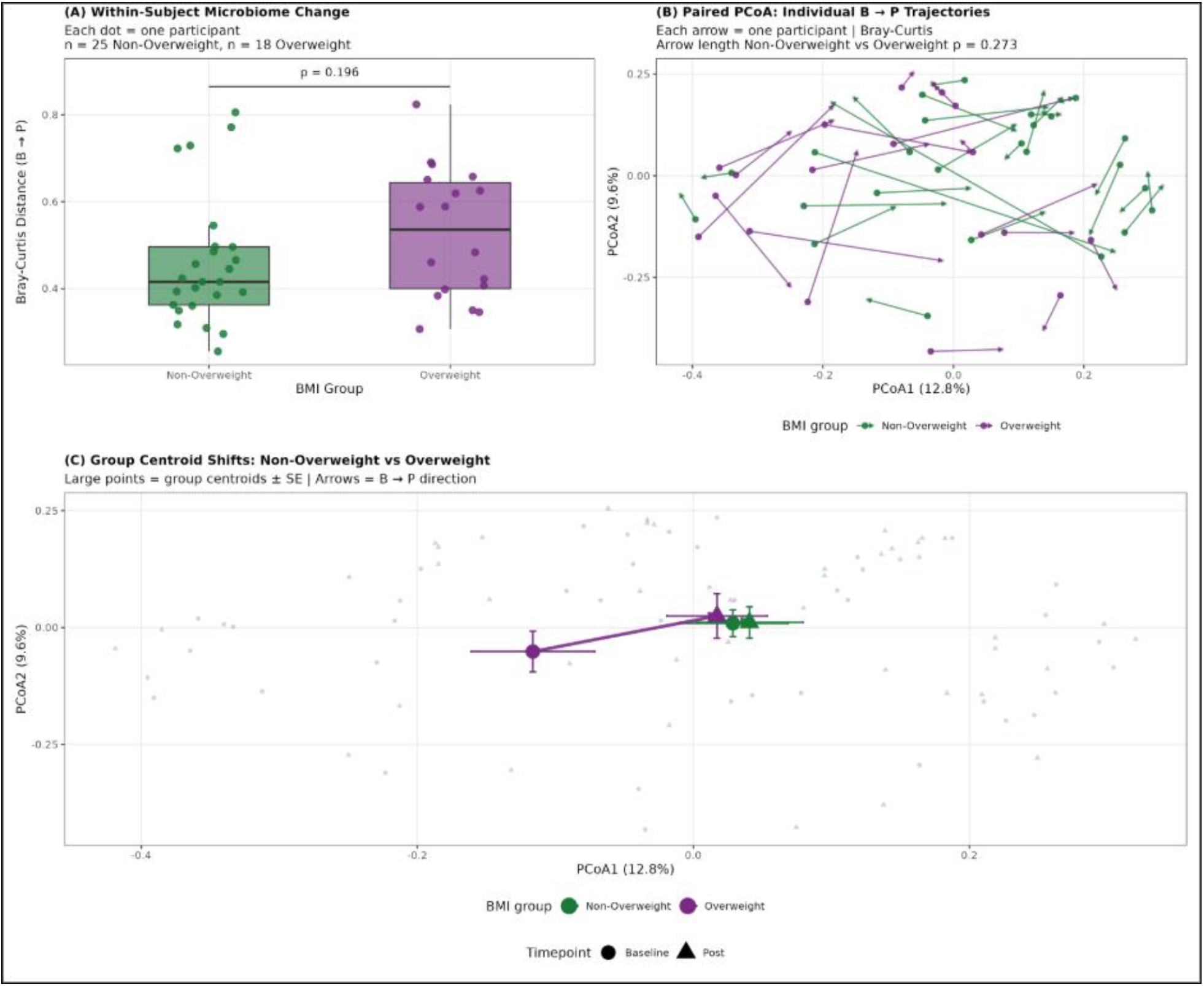
Three-panel comparison of within-subject change by BMI stratum. (A) Within-subject. B→P Bray-Curtis distance by stratum (Wilcoxon, p=0.196; mean 0.527 OW vs 0.460 Non-OW). **(B)** Paired-arrow PCoA connecting each participant’s baseline and post-intervention sample, with a permutation test on arrow length. **(C)** Group centroid shift from baseline to post-intervention (0.154 OW vs 0.012 Non-OW; 12.8× difference).

This directional consistency was also evident at the level of individual taxa, although no taxon reached FDR significance within the OW stratum. *Megamonas funiformis*, the leading SIMPER contributor, changed in the same direction in 73% of OW participants, and *Klebsiella michiganensis* was depleted in the majority of OW participants despite contributing little to bulk Bray-Curtis dissimilarity owing to its low absolute abundance **(Figure 6)**. The consistent per-participant direction of these changes supports the community-level centroid displacement described above.

**Fig. 6.**
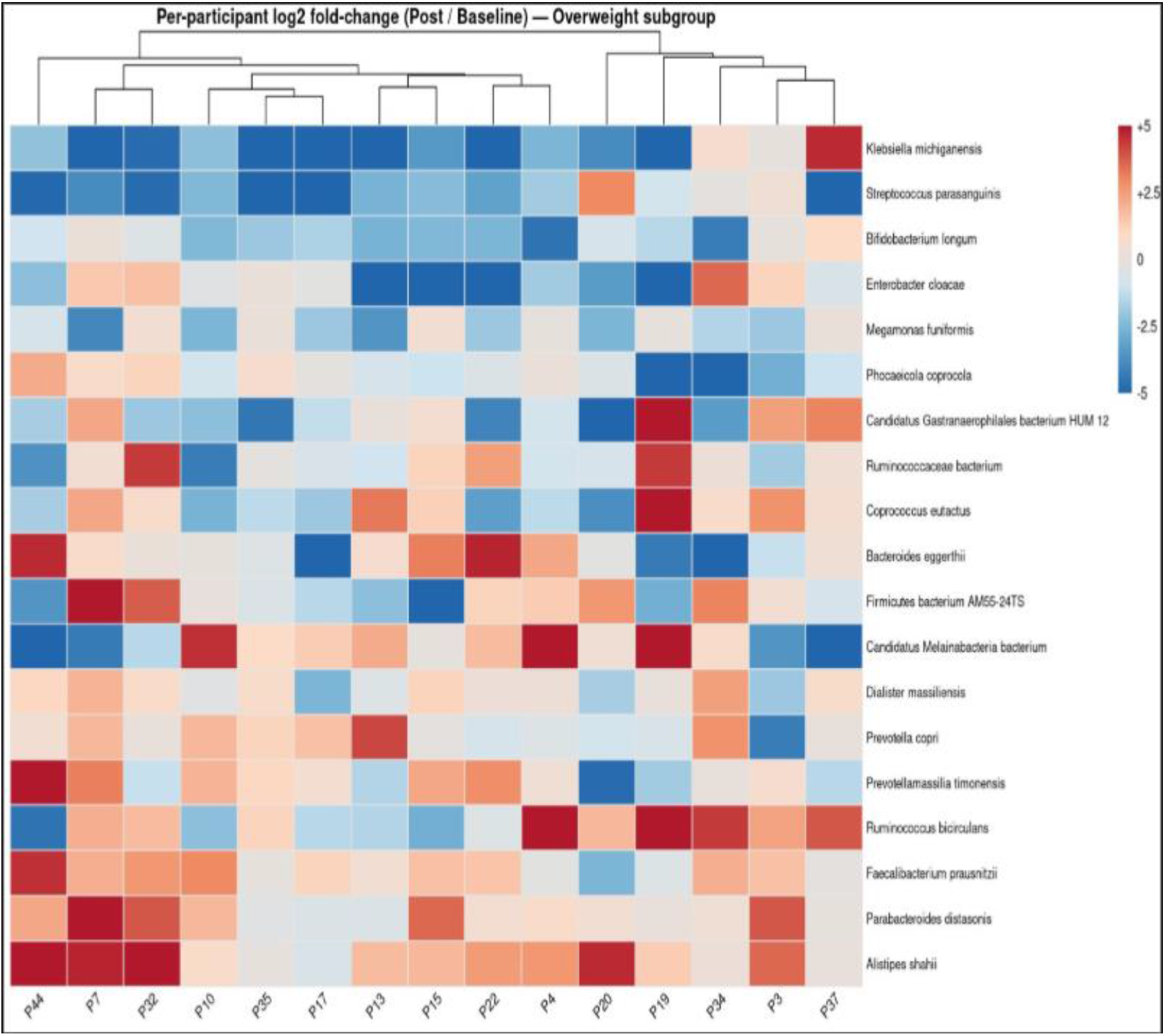
Per-participant direction of taxon-level change in overweight participants. Heatmap of per-participant log₂ fold-change (post ÷ baseline; pseudocount 10⁻⁶) for the union of the top 15 SIMPER contributors and all taxa reaching q < 0.25 in the OW-only MaAsLin2 model. Rows = taxa, ordered by mean fold-change; columns = individual OW participants (n = 18, hierarchically clustered). Blue = depletion, red = enrichment; values capped at ±5 log₂ units for display. No taxon shown reached FDR significance (q < 0.05) within the OW stratum; the panel is exploratory.

### 3.4 Depletion of oral-associated taxa and enrichment of butyrate producers characterize the compositional shift

To identify taxa responsible for the observed compositional change, differential abundance analysis was performed using MaAsLin2 with TSS-normalized abundance data, internal log transformation, and a linear mixed-effects model incorporating participant as a random intercept and timepoint, BMI group, region, and diet type as fixed effects. After filtering (prevalence ≥9 samples and mean relative abundance >10⁻⁴), 366 taxa were retained for analysis. Two species (*Streptococcus parasanguinis*, β = -2.33, q = 0.0102; *Gemella sanguinis*, β = -1.55, q = 0.021) reached FDR significance across the cohort. Both are oral-associated, and each was depleted after the intervention.

To increase statistical power for low-abundance taxa, differential abundance analysis was repeated at the genus level. Aggregation to the genus level yielded 312 genera, of which 140 entered testing after the same prevalence and abundance filters. Six genera remained significant after FDR correction **(Table 6, Supplementary Figure S3)**. Depletion of *Actinomyces* (q = 0.004), *Solobacterium* (q = 0.008), *Gemella* (q = 0.010), and *Streptococcus* (q = 0.013) and *Bifidobacterium* (q = 0.024), together with enrichment of *Faecalibacterium* (q = 0.016), confirmed that the dietary intervention affected broader taxonomic groups in addition to individual species. Notably, *Streptococcus*, *Gemella*, *Actinomyces*, and *Solobacterium* are all oral-origin genera with reported pathobiont behavior, and their coordinated depletion indicates that several oral-associated taxa changed in parallel following the intervention. The remaining two genera are probiotic. *Faecalibacterium* increased (q = 0.016); its species *F. prausnitzii* is a butyrate-producing commensal widely described as a next-generation probiotic (He et al., 2021). *Bifidobacterium* decreased (q = 0.024), with different species affected in the two BMI strata. The intervention therefore altered beneficial commensals as well as pathobionts, and not uniformly in the same direction. Genus-level aggregation strengthened several signals that were only suggestive at species level, particularly for *Actinomyces, Solobacterium,* and *Faecalibacterium*.

**Table 6.** Genus-level MaAsLin2 differential abundance (all-participants model); six genera FDR-significant. Coef = MaAsLin2 model coefficient (negative = depleted after the intervention, positive = increased post intervention); q = Benjamini-Hochberg adjusted p-value. The final column compares each genus-level result with the corresponding species-level result.

| Genus | Coef | q | Direction | Species-level comparison |
| --- | --- | --- | --- | --- |
| *Actinomyces* | -1.83 | 0.004 | ↓ | 12 species, none individually significant |
| *Solobacterium* | -1.32 | 0.008 | ↓ | strengthened from species $q = 0.086$ |
| *Gemella* | -1.31 | 0.010 | ↓ | Consistent (*G. sanguinis* $q = 0.021$) |
| *Streptococcus* | -1.50 | 0.013 | ↓ | 57 species, *S. parasanguinis* lead |
| *Faecalibacterium* | +0.79 | 0.016 | ↑ | *F. prausnitzii* species $q = 0.058 \rightarrow$ genus $q = 0.016$ |
| *Bifidobacterium* | -1.09 | 0.024 | ↓ | No individual species FDR-significant |

At the exploratory threshold (q < 0.25), 34 taxa were differentially abundant. Depleted taxa were predominantly oral-associated pathobiont organisms including *Enterococcus faecium*, whereas enriched taxa were mainly fibre-fermenting butyrate producers, including *Faecalibacterium prausnitzii* (+0.78, q = 0.058) and *Lachnospiraceae* bacterium GAM79 (+0.82, q = 0.153). Two beneficial species *Weissella cibaria* (−1.95, q = 0.052) and *Bifidobacterium adolescentis* (−1.29, q = 0.074), were also depleted. To assess the robustness, the whole-cohort analysis was repeated with LinDA and ALDEx2. Results were organized into three confidence tiers according to the number of tools in agreement. *Streptococcus parasanguinis* was the only Tier 1 signal, detected by all three methods and strongly supported by LinDA (padj = 0.0009). Approximately ten additional taxa formed Tier 2 (MaAsLin2 and LinDA), including *F. prausnitzii*, *B. adolescentis*, *W. cibaria*, *Megamonas funiformis*, *Oscillibacter*, *Phocaeicola* and *Parabacteroides distasonis*; the remainder were supported by MaAsLin2 alone. Results are provided in **Supplementary Table S5.**

Stratified MaAsLin2 models were then fitted separately within each BMI stratum to test whether the same taxa changed in both groups. No taxon reached FDR significance (q < 0.05) in either subgroup, which is expected given the reduced per-subgroup sample sizes; these models are therefore exploratory and are reported at q < 0.25. In OW participants (34 samples), five taxa showed exploratory changes: decreased *Streptococcus parasanguinis* (−2.34, q = 0.187), *Megamonas funiformis* (−1.51, q = 0.187), *Klebsiella michiganensis* (−3.26, q = 0.187) and *Bifidobacterium longum* (−1.58, q = 0.197), together with increased *Alistipes shahii* (+2.10, q = 0.187). In Non-OW participants (52 samples), seven taxa showed exploratory changes: depletion of *Solobacterium* sp. UBA3691 (−1.74, q = 0.080), *Roseburia* sp. AF12-17LB (−1.19, q = 0.165), *S. parasanguinis* (−2.15, q = 0.189), *Bifidobacterium adolescentis* (−1.61, q = 0.189), *Weissella cibaria* (−1.79, q = 0.192) and *Gemella sanguinis* (−1.36, q = 0.235), together with enrichment of *Faecalibacterium prausnitzii* (+0.62, q = 0.210). The genus *Bifidobacterium* split between subgroups, with *B. longum* depleted in OW and *B. adolescentis* depleted in Non-OW participants (**Supplementary Figure S4**).

To examine whether these compositional differences extended to the stable component of the community, core-microbiome membership was compared between timepoints within each BMI stratum. Turnover was 2.25-fold higher in OW than in Non-OW participants (Jaccard turnover 0.525 versus 0.233), consistent with the greater centroid displacement described above and providing a second, taxon-level line of evidence for BMI-dependent responsiveness. Notably, neither *S. parasanguinis* nor *G. sanguinis* was part of the core microbiome at either timepoint, despite being the two most robust differentially abundant species (**Supplementary Table S6, Supplementary Figure S5**).

### 3.5 Alpha diversity shows a trend from richness towards evenness in OW participants

We further tested whether the observed compositional shifts were accompanied by changes in within-sample microbial diversity by assessing six complementary alpha diversity metrics using per-participant linear mixed-effects models. Across the entire cohort, none of the metrics remained significant after false discovery rate (FDR) correction (smallest q = 0.119). Nevertheless, the direction of change was consistent across metrics, with all richness and phylogenetic diversity-based measures showing decreases, whereas the evenness-based measures shifted toward greater evenness. Stratification by BMI revealed that these trends were driven primarily by OW participants (**Figure 7**).

**Fig. 7.**
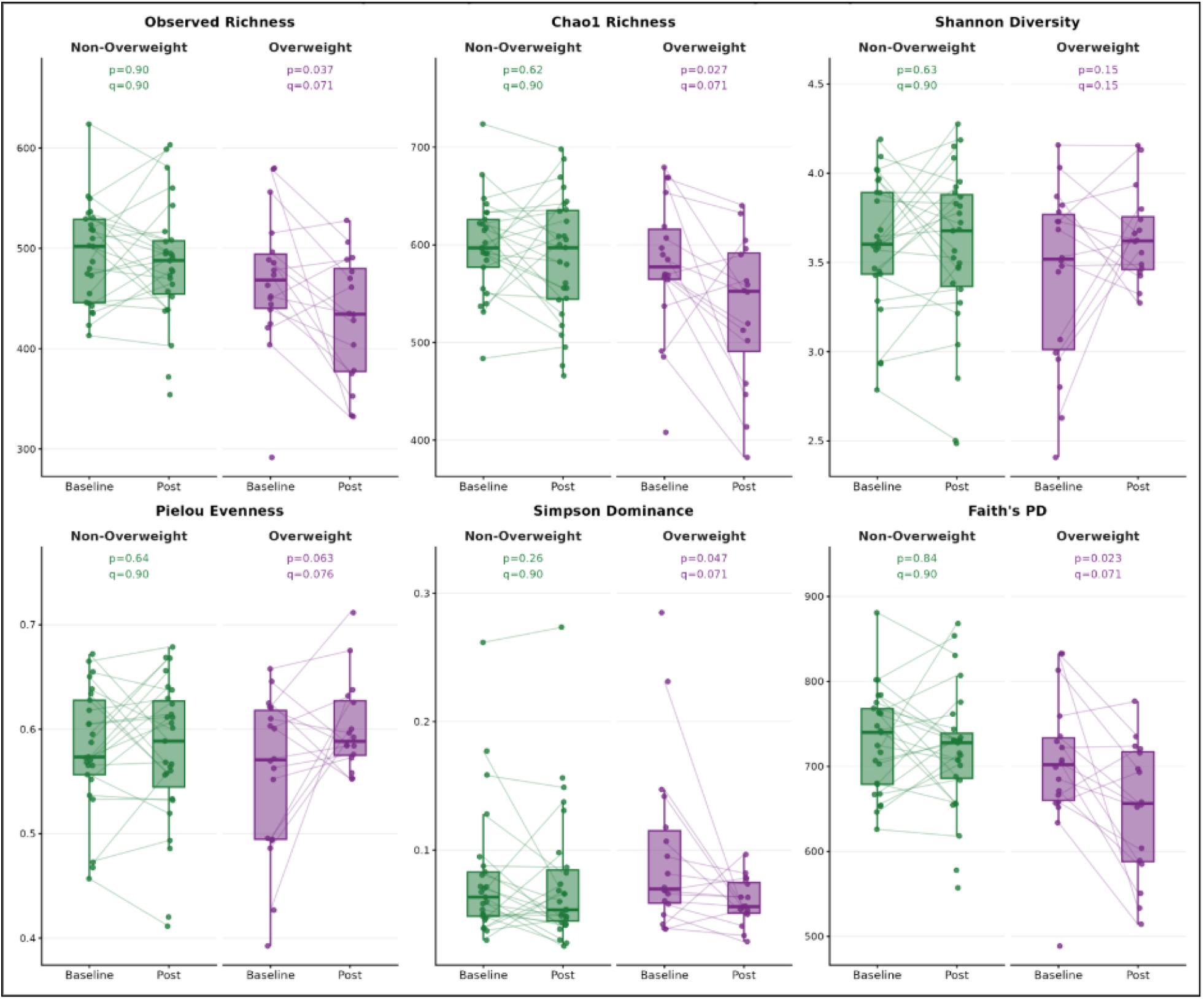
Paired baseline and post-intervention alpha diversity across six metrics. (Observed richness, Chao1, Shannon, Simpson, Pielou’s J, Faith’s PD), stratified by BMI stratum. Within each stratum, each metric was modelled with a linear mixed-effects model (nlme::lme) with timepoint as a fixed effect and a participant-level random intercept. Marginal and conditional R² were obtained with MuMIn.

Within the OW group, all six metrics moved in a consistent direction. The three richness indices declined nominally from baseline to post (Faith’s PD, p = 0.023; Chao1, p = 0.027; observed richness, p = 0.037). Simpson dominance fell (p = 0.047), and both Pielou’s evenness (p = 0.063) and Shannon diversity (p = 0.148) rose, all three indicating a shift toward greater evenness. None survived false-discovery-rate correction (all q ≈ 0.071-0.148). No index changed in the Non-OW group (lowest q = 0.897; **Supplementary Table S7**). Although not statistically significant after multiple-testing adjustments, the coordinated direction of these changes across all six metrics is consistent with a modest reshaping of the responsive subgroup toward fewer, more evenly distributed taxa. We therefore interpret this richness-to-evenness pattern as a trend-level, hypothesis-generating observation rather than a confirmed effect.

### 3.6 Functional potential is largely stable at whole-cohort level despite compositional change

Following the observed taxonomic restructuring, we next examined whether these changes were accompanied by alterations in microbial functional potential. Functional profiles were reconstructed from 154 KEGG orthologs (KOs) assigned to six metabolic pathways representing short-chain fatty acid production (acetate, butyrate, and propionate) and gas production (ammonia, methane, and hydrogen sulfide). Functional responses were assessed at the community, pathway, and KO levels.

Thirteen samples had fewer than 20 non-zero KOs and were identified as technically sparse. Whole-cohort functional tests were therefore repeated after excluding these samples, and additionally under CLR transformation, as a sensitivity check (**Supplementary Table S8**). At the whole-cohort level, functional beta-diversity showed no significant change following the intervention (p = 0.178-0.378), in contrast to the significant taxonomic shift observed in the same participants (p = 0.001), and neither the Timepoint × BMI (p = 0.154-0.668) nor the Timepoint × Region (p = 0.067-0.961) interaction was significant for functional composition. When examined by BMI group, no significant functional shift was observed in Non-OW participants, whereas a significant shift was detected in OW participants under CLR transformation (p = 0.01, R² = 10.1%). Betadisper on the sparse-sample-filtered functional dataset returned p = 0.002 for BMI group, indicating that functional dispersion differed between the two BMI strata. Because PERMANOVA is sensitive to differences in both group centroid and group dispersion, the OW-specific result may reflect this dispersion difference in addition to, or instead of, a centroid shift. Together with the smaller subgroup size (34 samples) and the exploratory nature of the stratified analysis, this finding is therefore reported as suggestive rather than confirmatory. In contrast, baseline functional composition was structured by geographic region under both the sparse-filtered TSS and CLR analyses (both p = 0.001) and by BMI under CLR transformation (p = 0.043), the same two factors that structured baseline taxonomic composition (Section 3.1), indicating that the functional dataset retained sufficient sensitivity to resolve between-participant differences despite the absence of a detectable timepoint effect.

At the pathway level, all six pathways showed a positive median change following the intervention. However, only methane showed a statistically significant increase after FDR correction (q = 0.041), while acetate, butyrate, propionate, and hydrogen sulfide showed suggestive increases (q = 0.057-0.083). Ammonia showed no significant change (q = 0.121) **(Table 7**, **Figure 8)**. Pathway-level Timepoint × BMI interactions, fitted as linear mixed-effects models on per-sample pathway sums, were null for all six pathways (all q > 0.93). Per-sample summed pathway abundances correlated strongly with sequencing depth (Spearman ρ > 0.80 for butyrate, acetate, propionate, and methane); pathway-level results are therefore reported alongside the KO-level analysis, which operates on within-pathway TSS-normalized abundances.

**Table 7.** Pathway-level paired Wilcoxon tests on per-sample summed KO abundances (n = 43 paired participants). Each row is a separate Wilcoxon signed-rank test on one pathway. V = signed-rank statistic; P = unadjusted p-value; q = Benjamini-Hochberg adjusted p-value; Median = median per-participant difference (post-baseline).

| Pathway | V | p | q | Median | Significance |
| --- | --- | --- | --- | --- | --- |
| Methane | 694 | 0.0069 | 0.041 | +261 | FDR-significant |
| Acetate | 654 | 0.028 | 0.057 | +574 | FDR-suggestive |
| Butyrate | 666 | 0.019 | 0.057 | +819 | FDR-suggestive |
| Propionate | 624 | 0.069 | 0.083 | +360 | FDR-suggestive |
| Hydrogen sulfide | 575.5 | 0.061 | 0.083 | +30 | FDR-suggestive |
| Ammonia | 576 | 0.121 | 0.121 | +393 | n.s. |

**Fig. 8.**
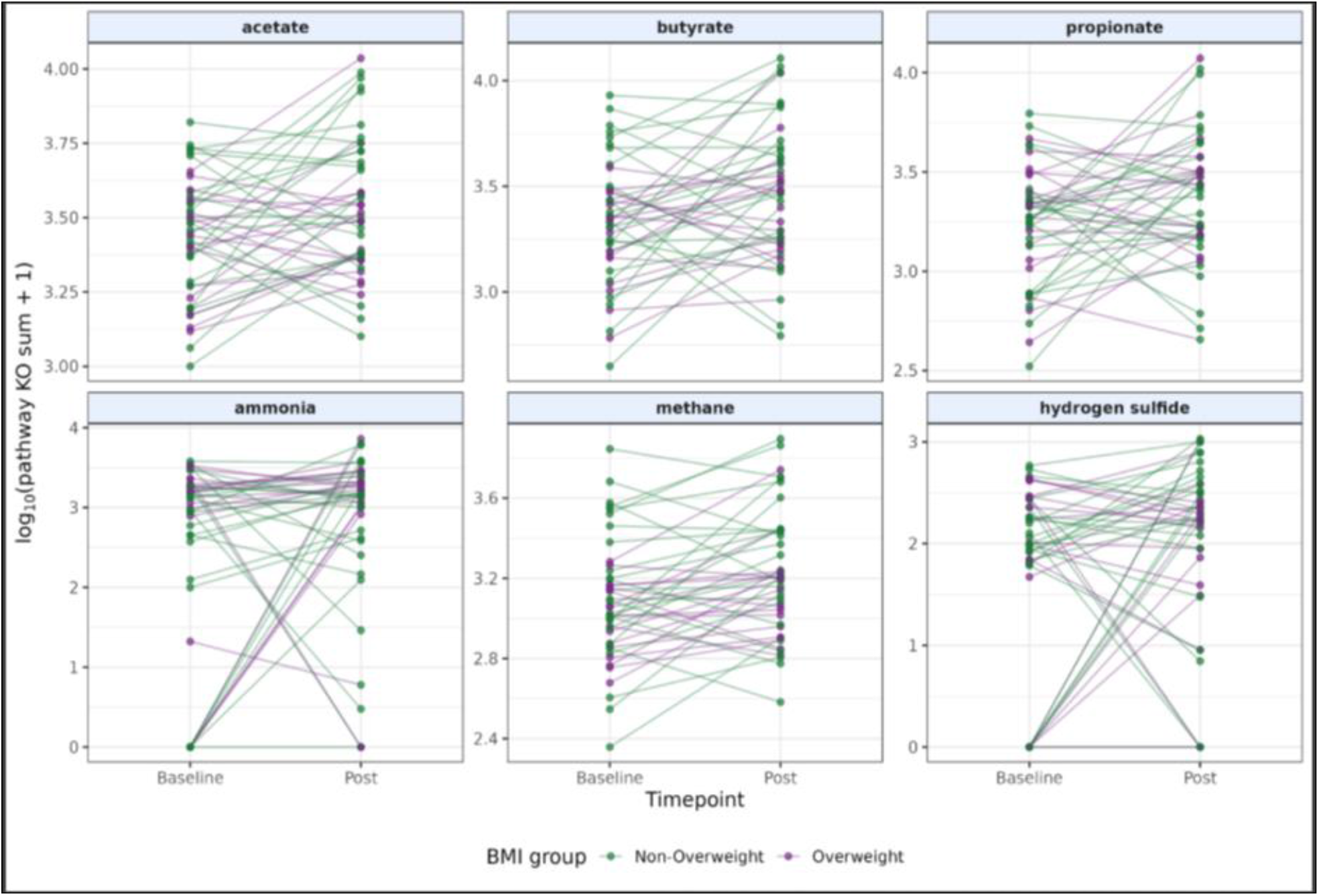
Pathway-level functional response to intervention. Paired baseline (B) and post-intervention (P) pathway-level KO abundances for acetate, butyrate, propionate, ammonia, methane, and hydrogen sulfide (H₂S). Each line connects paired measurements from the same participant, with BMI groups shown in green (Non-OW) and purple (OW).

At the KO level, two KOs showed FDR-significant changes across the whole cohort, both within the hydrogen sulfide production pathway (K05396, β = −0.89, q = 0.045; K20021, β = +0.48, q = 0.045). BMI-stratified analysis identified more differentially abundant KOs in OW than in Non-OW participants (34 versus 4 at q < 0.25). The strongest OW signals were in the ammonia pathway (K10672, − 0.89, q < 0.001) and the butyrate pathway (K01844, +1.67, q = 0.007). These exploratory findings are provided in **Supplementary Table S9** and **Supplementary Figure S6.**

We also performed a paired-swap permutation analysis to further assess the consistency of the observed pathway-level changes. The observed 6/6-positive pattern occurred in 107/1,000 permutations in the whole cohort (empirical p = 0.107) and in 154/1,000 permutations in Non-OW participants (p = 0.154). In OW participants, the observed pattern of five positive pathways and a negative hydrogen sulfide pathway occurred in 61/1,000 permutations (p = 0.061). Thus, the overall pathway-level directional pattern was suggestive but not definitive; detailed results are provided in **Supplementary Table S10** and **Supplementary Figure S7**.

## 4. Discussion

Dietary patterns rich in whole grains, vegetables, fibre, and fermented foods are widely associated with favorable gut microbial and metabolic characteristics (Leeming et al., 2019). However, heterogeneity in dietary exposures, study populations, and environmental conditions has limited the ability to define consistent microbiome responses across cohorts (Falony et al., 2016; Rothschild et al., 2018). To minimize these sources of variability, the present study was conducted in a pop-up village with relatively similar environmental and lifestyle characteristics, allowing the determinants of microbial responsiveness to be examined more directly. Our findings demonstrate that response to the short-term dietary intervention varied according to baseline host characteristics, with a more pronounced community-level taxonomic response observed among OW participants than among Non-OW participants. This observation is consistent with previous evidence showing that microbiome responses to dietary changes can be highly individualized and influenced by baseline microbial characteristics (Johnson et al., 2019).

At baseline, geographic region and BMI were significantly associated with gut microbial composition in our cohort. Geographic differences in gut microbiome composition have been reported previously, reflecting variation between populations and environments (Deschasaux et al., 2018; Gupta et al., 2017; Yatsunenko et al., 2012). Although the geographic region structured the community at baseline, it did not modify the community-level response to the intervention. The response differed instead by BMI group, with a significant shift in OW participants but not in Non-OW participants. This pattern is consistent with previous studies indicating that variation in host and microbial characteristics may contribute to differences in responses to dietary interventions (Fassarella et al., 2021; Klimenko et al., 2022), although the present findings are specific to our cohort and do not establish BMI as a general determinant of microbiome responsiveness. Habitual diet type was not associated with baseline community composition (p = 0.792), nor with the response to the intervention (Timepoint × Diet, p = 0.483). This differs from studies reporting differences between vegetarian and omnivorous eating patterns. Those studies, however, examined individual taxa rather than whole-community structure (De Filippis et al., 2016; Kabeerdoss et al., 2012), and Indian comparisons have drawn diet groups from separate regions, so diet and geography cannot be distinguished (Dhakan et al., 2019). When the same composition-level test was applied to 21,561 individuals, diet pattern accounted for only 0.2-2.8% of variance in beta diversity (Fackelmann et al., 2025). A between-subject effect of this magnitude is not detectable in 25 versus 18 participants, whereas the timepoint effect reported here was tested within participants, where inter-individual variation does not contribute to the comparison. Diet type in our study was also recorded as a single self-reported binary label rather than as quantified intake. This null therefore reflects our sample size and how diet was measured and should not be taken as evidence that diet type has no effect.

The intervention was associated with a modest but consistent shift in microbial community composition across the cohort, indicating that the gut microbiome was responsive to the dietary change over the 15-day intervention period. The relatively small magnitude of the shift is consistent with the fact that dietary exposure represents only one component of the multiple factors shaping gut microbial communities (Gacesa et al., 2022; Zhernakova et al., 2016), while individual dietary factors typically explain a limited proportion of population-level microbiome variation (Falony et al., 2016). The short duration of the intervention is also important when considering the temporal nature of the observed response. Although the gut microbiome can respond rapidly to dietary perturbations, our sampling captured only the immediate post-intervention state. Because no washout or subsequent follow-up sample was collected, we cannot determine whether the observed community-level changes were sustained beyond the 15-day intervention or subsequently moved towards baseline. Similarly, shorter intervention periods may initiate microbial transitions without allowing complete ecological restructuring (Leeming et al., 2019). Therefore, the coordinated community shift observed in our study represents a measurable response during the 15-day intervention period, while whether longer-term dietary exposure would result in more sustained microbial changes remains to be determined.

The significant shift in microbial composition was accompanied by stable beta-dispersion across all distance metrics, indicating that the intervention did not increase variability between participants. Rather than producing divergent individual trajectories, the intervention shifted the overall microbial community centroid while maintaining comparable within-group dispersion. This indicates a coordinated community-level response rather than increased ecological instability. Compared with the extreme dietary crossover reported by David et al. (2014), where complete animal-versus plant-based dietary replacement produced rapid and pronounced microbiome restructuring, the present intervention represented a more moderate, real-world dietary exposure. Therefore, the smaller effect size observed here is expected, and the reproducibility of the shift across multiple beta-diversity metrics suggests that consistency of response, rather than magnitude alone, is an important indicator of biological relevance in short-term dietary interventions.

Importantly, the response was not uniform across participants. The dietary shift was primarily observed in OW individuals, whereas Non-OW participants showed limited change. This difference did not arise from larger individual changes in OW participants, whose within-subject dissimilarities were comparable to those of Non-OW participants (p = 0.196). Instead, it arose from greater consistency in the direction of change across individuals, reflected in a substantially larger displacement of the group centroid in OW participants and in the concordant per-participant direction of change for individual taxa. This indicates that BMI influenced the extent to which the microbiome responded to dietary intervention. Individuals with higher adiposity have previously been shown to harbor distinct microbial configuration compared with Non-OW individuals, including differences in microbial richness and community structure that may contribute to variation in responses to dietary perturbations (Cotillard et al., 2013; Le Chatelier et al., 2013). Consistent with this concept, a recent study has demonstrated that baseline microbiome characteristics can determine whether individuals exhibit measurable microbiome responses to dietary interventions (Hoffmann Sardá et al., 2025; Klimenko et al., 2022). The limited response observed in Non-OW participants may therefore reflect differences in baseline community configuration or ecological stability rather than an absence of biological responsiveness. Such inter-individual variability supports the emerging concept of precision nutrition, where host characteristics and baseline microbiome features influence the effectiveness of dietary interventions (Berry et al., 2020; Wastyk et al., 2021; Zeevi et al., 2015). Within the OW subgroup, the observed alpha-diversity pattern, characterized by reduced richness and phylogenetic diversity together with increased evenness, suggests ecological restructuring of community organization rather than a simple loss of microbial complexity. Changes in diversity metrics should therefore be interpreted in the context of accompanying compositional shifts and ecological function, as reductions in richness do not necessarily indicate impaired microbial health when community balance and functional capacity are maintained (Valdes et al., 2018). Although these associations did not survive FDR correction, the consistent direction of change across multiple diversity metrics suggests a trend-level ecological restructuring rather than a confirmed alteration in microbial diversity. Similar reductions in microbial richness and phylogenetic diversity have been reported in adults with high adiposity and impaired metabolic health, including associations between low microbial gene richness, adiposity, insulin resistance, and inflammation (Le Chatelier et al., 2013; Kim et al., 2020). The concurrent tendency toward increased evenness may represent a short-term reorganization of community structure following increased fibre availability; however, this interpretation remains hypothesis-generating and requires validation in larger cohorts.

At the level of individual taxa, the depletion of the oral microbe *Streptococcus parasanguinis* represented the most consistent signal across differential-abundance approaches. Although typically considered commensal in oral cavity, it may exhibit pathobiont-like behavior under specific ecological conditions. Pathobionts are members of the resident microbiota that are normally tolerated by the host within their native ecological niche but can contribute to disease-associated states when environmental or host factors promote their expansion or altered activity, such as immune dysfunction or microbial dysbiosis (Chow et al., 2011). The concurrent depletion of *Gemella sanguinis*, together with the absence of both taxa from the core microbiome at both timepoints, suggests that these organisms were transient members of the gut community rather than stable resident taxa in this cohort. The depletion of *Klebsiella michiganensis* in the OW subgroup is consistent with this pattern, as this taxon belongs to a group that has been associated with pathobiont-like behavior and dysbiotic states under specific ecological conditions (Atarashi et al., 2017; Shin et al., 2015). *Actinomyces* was depleted at the genus level (q = 0.004), although no individual species reached significance. These members of the resident microbiota may contribute to inflammatory states when ecological disruption, increased abundance, or altered host-microbe interactions favour their expansion or activity (Chow et al., 2011). Therefore, their reduction following a fibre-rich, minimally processed dietary intervention is consistent with a restructuring of the microbial community away from taxa associated with dysbiotic states and towards a more balanced ecological profile.

In parallel, fibre-fermenting butyrate-producing genus *Faecalibacterium* was enriched, driven by *Faecalibacterium prausnitzii,* a beneficial commensal with anti-inflammatory and gut-barrier supporting properties (Sokol et al., 2008). Two further butyrate producers, *Roseburia hominis* and *Agathobaculum butyriciproducens*, also entered the OW-subgroup core microbiome following the intervention. The loss of butyrate producers such as *R. hominis* is itself a recognised feature of intestinal dysbiosis (Machiels et al., 2014), so their enrichment reflects a shift toward a healthier community state. This is consistent with increased fermentable fibre availability, as fibre-utilising microbes convert dietary substrates into short-chain fatty acids (SCFAs), particularly butyrate, which contributes to epithelial barrier maintenance, mucosal immune regulation, and host metabolic health (Blaak et al., 2020; Koh et al., 2016; Makki et al., 2018; Parada Venegas et al., 2019; Zhang et al., 2023). Similar increases in SCFA-associated microbial taxa have been reported following plant-rich and Mediterranean-style dietary patterns, which provide greater amounts of fermentable fibre (De Filippis et al., 2016; Ghosh et al., 2020; Nagpal et al., 2019;). On the other hand, *Bifidobacterium* showed contrasting species-level responses between subgroups, with *Bifidobacterium longum* depleted in OW participants and *Bifidobacterium adolescentis* depleted in Non-OW participants. This finding highlights that microbial responses to the same dietary intervention may differ according to the host metabolic context and that genus-level interpretations may overlook important species-specific variation (Milani et al., 2016).

In contrast to the taxonomic changes, the overall functional profile remained relatively stable at the whole-cohort level. At the pathway level, methane was the only pathway to reach FDR significance (q = 0.041), but per-sample summed pathway abundances correlated strongly with sequencing depth (Spearman ρ > 0.80), so this result is not interpreted as a biological effect. Two KOs within the hydrogen sulfide pathway reached FDR significance at the whole-cohort level (K05396, K20021; q = 0.045). Several pathways showed directional changes, particularly in the OW subgroup, including an increase in the butyrate-related function K01844 (q = 0.007) and decreases in several ammonia-assimilation functions (q < 0.001) within the BMI-stratified models. These stratified analyses were exploratory, were not corrected across subgroups, and did not reach significance in the whole-cohort model. They should therefore be interpreted as hypothesis-generating rather than confirmatory. The overall pattern of taxonomic change alongside limited FDR-adjusted functional changes is compatible with functional redundancy, whereby different microbial taxa may contribute to similar metabolic functions and maintain overall functional potential despite changes in community composition (Moya & Ferrer, 2016). However, functional redundancy cannot be established directly from the present data, which assesses predicted functional potential rather than measured microbial activity.

Several limitations should be considered when evaluating these findings. First, the relatively small sample size (n = 43) and modest community-level effect sizes (whole-cohort R² = 1.15-1.58%) limited statistical power, particularly for subgroup analyses and after correction for multiple comparisons. Several alpha-diversity measures and the exploratory functional permutation analysis in the OW subgroup showed borderline statistical evidence and require validation in larger cohorts. Second, 15 days intervention period was relatively short compared with longer-term dietary intervention studies (e.g., Ghosh et al., 2020), and microbiome profiling was limited to baseline and post-intervention time points. Therefore, the study could not determine whether the observed microbiome changes would persist or return toward baseline or characterize the rate and progression of these changes over time. Additional longitudinal sampling would be required to assess the temporal stability and reversibility of these changes. Third, individual-level dietary intake was not quantitatively recorded, so dose–response relationships between specific dietary components and microbiome features cannot be established. The intervention is treated throughout as a shared dietary environment rather than a per-participant standardized exposure. Fourth, the pop-up village design limits the ability to distinguish the effects of the shared diet from those of the shared living environment. Cohabitation may influence gut microbiome composition through microbial transmission and shared environmental exposures. Therefore, some of the observed microbiome changes may be attributable to cohabitation in addition to dietary intervention. The study was also single-arm, with no concurrent non-intervention control group, so the observed changes cannot be separated from study-participation effects. Fifth, Oxford Nanopore metagenomic sequencing may have reduced sensitivity for some low-abundance taxa compared with Illumina sequencing, although taxonomic abundance estimates were comparable for high-abundance species (Wongsurawat et al., 2019). To minimize the influence of sparsely detected features, association testing was restricted to features with a prevalence of ≥9 samples and a mean relative abundance >10⁻⁴.

Sixth, BMI stratification was observational, and the stronger microbiome response observed among overweight participants cannot be attributed to BMI alone. Because both baseline BMI (p = 0.024) and baseline geographic region (p = 0.046) were associated with community composition, the observed response differences may reflect baseline microbiome characteristics, BMI, geographic background, or their joint contribution. Seventh, functional analyses were based on gene-content-derived functional potential rather than direct measurements of microbial activity or metabolite production. Therefore, the observed changes in functional potential cannot be taken as direct evidence of changes in microbial pathway activity and require validation using metatranscriptomics and metabolomics approaches. Eighth, the binary BMI classification (<25 versus ≥25 kg/m²) was adopted because of the limited sample size. This classification limited the assessment of microbiome responsiveness across the full BMI range and prevented evaluation of category-specific response patterns among underweight, normal-weight, overweight, and obesity categories. Finally, the open recruitment approach was a limitation, as participation was voluntary, and the cohort was not recruited according to predefined clinical or metabolic categories. Consequently, specific clinical or metabolic subgroups were not systematically represented, which may limit the generalizability of the findings across different health states.

## 5. Conclusion

This study shows that a short-term, gut-friendly dietary intervention conducted in a controlled pop-up village setting was associated with a modest but consistent shift in gut microbial community composition. OW participants showed a greater community-level compositional response, together with a suggestive shift from richness to evenness-based community organization, whereas whole-cohort differential-abundance analyses identified depletion of several oral-associated taxa and enrichment of fibre-fermenting taxa. In contrast, Non-OW participants showed comparatively limited alterations in microbial composition. Despite these taxonomic changes, the predicted gene content of six targeted metabolic pathways remained largely stable at the whole-cohort level. This is consistent with functional redundancy.

These findings support an emerging view that microbiome responsiveness to diet depends on host-microbiome characteristics, rather than following a one-size-fits-all pattern. Future studies should incorporate longer intervention and follow-up periods, quantitative dietary assessment, metabolomic and metatranscriptomic validation, and cohorts spanning the full BMI spectrum, to define the host and microbial features that predict responsiveness and to inform the design of individualized nutritional strategies.

## Supporting information

Metadata

## Acknowledgments

The authors thank ResearchHub, the volunteers of ZuGrama, and the participants of the study. We are grateful to the Hyatt Food and Beverage team for their support with meal provision throughout the study, and to Dr. Suramya Aasthana (Indian Institute of Science, Bangalore) for valuable input.

## Author Contributions

D.P. conceived and designed the study. A.M. performed laboratory work. L.J. performed bioinformatics and statistical analysis. S.U. and L.J. wrote the original draft. O.M. and D.P. reviewed and edited the manuscript. All authors read and approved the final manuscript.

## Informed Consent Statement

Informed consent was obtained from all subjects involved in the study.

## Data Availability Statement

De-identified summary data and analysis code are available from the corresponding author upon request.

## References

Abdill, R. J., Adamowicz, E. M., & Blekhman, R. (2022). Public human microbiome data are dominated by highly developed countries. PLoS biology, 20(2), e3001536.

Atarashi, K., Suda, W., Luo, C., Kawaguchi, T., Motoo, I., Narushima, S., … & Honda, K. (2017). Ectopic colonization of oral bacteria in the intestine drives TH1 cell induction and inflammation. Science, 358(6361), 359–365.

Bartoń, K. 2023. MuMIn: multi-model inference. (R package).

Beckman Coulter, 2016. Agencourt AMPure XP: PCR purification (product insert). Brea, CA: Beckman Coulter, Inc.

Berry, S. E., Valdes, A. M., Drew, D. A., Asnicar, F., Mazidi, M., Wolf, J., … & Spector, T. D. (2020). Human postprandial responses to food and potential for precision nutrition. Nature medicine, 26(6), 964–973.

Blaak, E. E., Canfora, E. E., Theis, S., Frost, G., Groen, A. K., Mithieux, G., … & Verbeke, K. (2020). Short chain fatty acids in human gut and metabolic health. Beneficial microbes, 11(5), 411–455.

Chow, J., Tang, H., & Mazmanian, S. K. (2011). Pathobionts of the gastrointestinal microbiota and inflammatory disease. Current opinion in immunology, 23(4), 473–480.

Clemente-Suárez, V. J., Beltrán-Velasco, A. I., Redondo-Flórez, L., Martín-Rodríguez, A., & Tornero-Aguilera, J. F. (2023). Global impacts of western diet and its effects on metabolism and health: a narrative review. Nutrients, 15(12), 2749.

Corbin, K. D., Carnero, E. A., Dirks, B., Igudesman, D., Yi, F., Marcus, A., … & Smith, S. R. (2023). Host-diet-gut microbiome interactions influence human energy balance: a randomized clinical trial. Nature communications, 14(1), 3161.

Cotillard, A., Kennedy, S. P., Kong, L. C., Prifti, E., Pons, N., Le Chatelier, E., … & Ehrlich, S. D. (2013). Dietary intervention impact on gut microbial gene richness. Nature, 500(7464), 585–588.

David, L. A., Maurice, C. F., Carmody, R. N., Gootenberg, D. B., Button, J. E., Wolfe, B. E., … & Turnbaugh, P. J. (2014). Diet rapidly and reproducibly alters the human gut microbiome. Nature, 505(7484), 559–563.

De Filippis, F., Pellegrini, N., Vannini, L., Jeffery, I. B., La Storia, A., Laghi, L., … & Ercolini, D. (2016). High-level adherence to a Mediterranean diet beneficially impacts the gut microbiota and associated metabolome. Gut, 65(11), 1812–1821.

Deschasaux, M., Bouter, K. E., Prodan, A., Levin, E., Groen, A. K., Herrema, H., … & Nieuwdorp, M. (2018). Depicting the composition of gut microbiota in a population with varied ethnic origins but shared geography. Nature medicine, 24(10), 1526–1531.

Dhakan, D. B., Maji, A., Sharma, A. K., Saxena, R., Pulikkan, J., Grace, T., … & Sharma, V. K. (2019). The unique composition of Indian gut microbiome, gene catalogue, and associated fecal metabolome deciphered using multi-omics approaches. Gigascience, 8(3), giz004.

Dubey, A. K., Uppadhyaya, N., Nilawe, P., Chauhan, N., Kumar, S., Gupta, U. A., & Bhaduri, A. (2018). LogMPIE, pan-India profiling of the human gut microbiome using 16S rRNA sequencing. Scientific data, 5(1), 180232.

Fackelmann, G., Manghi, P., Carlino, N., Heidrich, V., Piccinno, G., Ricci, L., … & Segata, N. (2025). Gut microbiome signatures of vegan, vegetarian and omnivore diets and associated health outcomes across 21,561 individuals. Nature microbiology, 10(1), 41–52.

Falony, G., Joossens, M., Vieira-Silva, S., Wang, J., Darzi, Y., Faust, K., … & Raes, J. (2016). Population-level analysis of gut microbiome variation. Science, 352(6285), 560–564.

Fassarella, M., Blaak, E. E., Penders, J., Nauta, A., Smidt, H., & Zoetendal, E. G. (2021). Gut microbiome stability and resilience: elucidating the response to perturbations in order to modulate gut health. Gut, 70(3), 595–605.

Fernández-Pato, A., Sinha, T., Gacesa, R., Andreu-Sánchez, S., Gois, M. F. B., Gelderloos-Arends, J., … & Kurilshikov, A. (2024). Choice of DNA extraction method affects stool microbiome recovery and subsequent phenotypic association analyses. Scientific Reports, 14(1), 3911.

Gacesa, R., Kurilshikov, A., Vich Vila, A., Sinha, T., Klaassen, M. A., Bolte, L. A., … & Weersma, R. K. (2022). Environmental factors shaping the gut microbiome in a Dutch population. Nature, 604(7907), 732–739.

Ghosh, T. S., Rampelli, S., Jeffery, I. B., Santoro, A., Neto, M., Capri, M., … & O’Toole, P. W. (2020). Mediterranean diet intervention alters the gut microbiome in older people reducing frailty and improving health status: the NU-AGE 1-year dietary intervention across five European countries. Gut, 69(7), 1218–1228.

Gupta, V. K., Paul, S., & Dutta, C. (2017). Geography, ethnicity or subsistence-specific variations in human microbiome composition and diversity. Frontiers in microbiology, 8, 1162.

He, X., Zhao, S., & Li, Y. (2021). Faecalibacterium prausnitzii: A next-generation probiotic in gut disease improvement. Canadian Journal of Infectious Diseases and Medical Microbiology, 2021(1), 6666114.

He, Y., Wang, B., Wen, L., Wang, F., Yu, H., Chen, D., … & Zhang, C. (2022). Effects of dietary fiber on human health. Food Science and Human Wellness, 11(1), 1–10.

Hoffmann Sardá, F. A., Giuntini, E. B., Oliveira, A., Souza, G. S., Prado, S. B., Taddei, C. R., … & Hoffmann, C. (2025). Baseline intestinal microbiota composition influences response to a real-world dietary fiber intervention. npj Biofilms and Microbiomes, 11(1), 203.

Holscher, H. D. (2017). Dietary fiber and prebiotics and the gastrointestinal microbiota. Gut microbes, 8(2), 172–184.

Jackson, M. A., Verdi, S., Maxan, M. E., Shin, C. M., Zierer, J., Bowyer, R. C., … & Steves, C. J. (2018). Gut microbiota associations with common diseases and prescription medications in a population-based cohort. Nature communications, 9(1), 2655.

Johnson, A. J., Vangay, P., Al-Ghalith, G. A., Hillmann, B. M., Ward, T. L., Shields-Cutler, R. R., … & Knights, D. (2019). Daily sampling reveals personalized diet-microbiome associations in humans. Cell host & microbe, 25(6), 789–802.

Kabeerdoss, J., Devi, R. S., Mary, R. R., & Ramakrishna, B. S. (2012). Faecal microbiota composition in vegetarians: comparison with omnivores in a cohort of young women in southern India. British Journal of Nutrition, 108(6), 953–957.

Kim, M. H., Yun, K. E., Kim, J., Park, E., Chang, Y., Ryu, S., … & Kim, H. N. (2020). Gut microbiota and metabolic health among overweight and obese individuals. Scientific reports, 10(1), 19417.

Klimenko, N. S., Odintsova, V. E., Revel-Muroz, A., & Tyakht, A. V. (2022). The hallmarks of dietary intervention-resilient gut microbiome. npj Biofilms and Microbiomes, 8(1), 77.

Koh, A., De Vadder, F., Kovatcheva-Datchary, P., & Bäckhed, F. (2016). From dietary fiber to host physiology: short-chain fatty acids as key bacterial metabolites. Cell, 165(6), 1332–1345.

Le Chatelier, E., Nielsen, T., Qin, J., Prifti, E., Hildebrand, F., Falony, G., … & Pedersen, O. (2013). Richness of human gut microbiome correlates with metabolic markers. Nature, 500(7464), 541–546.

Leeming, E. R., Johnson, A. J., Spector, T. D., & Le Roy, C. I. (2019). Effect of diet on the gut microbiota: rethinking intervention duration. Nutrients, 11(12), 2862.

Machiels, K., Joossens, M., Sabino, J., De Preter, V., Arijs, I., Eeckhaut, V., … & Vermeire, S. (2014). A decrease of the butyrate-producing species Roseburia hominis and Faecalibacterium prausnitzii defines dysbiosis in patients with ulcerative colitis. Gut, 63(8), 1275–1283.

Makki, K., Deehan, E. C., Walter, J., & Bäckhed, F. (2018). The impact of dietary fiber on gut microbiota in host health and disease. Cell host & microbe, 23(6), 705–715.

Marić, J., Križanović, K., Riondet, S., Nagarajan, N., & Šikić, M. (2024). Comparative analysis of metagenomic classifiers for long-read sequencing datasets. BMC bioinformatics, 25(1), 15.

Milani, C., Turroni, F., Duranti, S., Lugli, G. A., Mancabelli, L., Ferrario, C., … & Ventura, M. (2016). Genomics of the genus Bifidobacterium reveals species-specific adaptation to the glycan-rich gut environment. Applied and Environmental Microbiology, 82(4), 980–991.

Moya, A., & Ferrer, M. (2016). Functional redundancy-induced stability of gut microbiota subjected to disturbance. Trends in microbiology, 24(5), 402–413.

Nagpal, R., Neth, B. J., Wang, S., Craft, S., & Yadav, H. (2019). Modified Mediterranean-ketogenic diet modulates gut microbiome and short-chain fatty acids in association with Alzheimer’s disease markers in subjects with mild cognitive impairment. EBioMedicine, 47, 529–542.

Nakagawa, S. and Schielzeth, H. (2013), A general and simple method for obtaining *R*^2^ from generalized linear mixed-effects models. Methods Ecol Evol, 4: 133–142.

Oxford Nanopore Technologies, 2023. Native Barcoding Kit 24 V14 (SQK-NBD114.24): protocol and PromethION Flow Cell (R10.4.1) documentation. Oxford: Oxford Nanopore Technologies plc.

Parada Venegas, D., De la Fuente, M. K., Landskron, G., González, M. J., Quera, R., Dijkstra, G., … & Hermoso, M. A. (2019). Short chain fatty acids (SCFAs)-mediated gut epithelial and immune regulation and its relevance for inflammatory bowel diseases. Frontiers in immunology, 10, 277.

Qiagen, 2020. QIAamp Fast DNA Stool Mini Kit Handbook (cat. no. 51604). Hilden, Germany: Qiagen GmbH.

Rothschild, D., Weissbrod, O., Barkan, E., Kurilshikov, A., Korem, T., Zeevi, D., … & Segal, E. (2018). Environment dominates over host genetics in shaping human gut microbiota. Nature, 555(7695), 210–215.

Shin, N. R., Whon, T. W., & Bae, J. W. (2015). Proteobacteria: microbial signature of dysbiosis in gut microbiota. Trends in biotechnology, 33(9), 496–503.

Sokol, H., Pigneur, B., Watterlot, L., Lakhdari, O., Bermúdez-Humarán, L. G., Gratadoux, J. J., … & Langella, P. (2008). Fecalibacterium prausnitzii is an anti-inflammatory commensal bacterium identified by gut microbiota analysis of Crohn disease patients. Proceedings of the National Academy of Sciences, 105(43), 16731–16736.

Thermo Fisher Scientific, 2015. Qubit dsDNA HS Assay Kit (product information sheet). Waltham, MA: Thermo Fisher Scientific Inc.

Valdes, A. M., Walter, J., Segal, E., & Spector, T. D. (2018). Role of the gut microbiota in nutrition and health. Bmj, 361.

Valles-Colomer, M., Blanco-Míguez, A., Manghi, P., Asnicar, F., Dubois, L., Golzato, D., … & Segata, N. (2023). The person-to-person transmission landscape of the gut and oral microbiomes. Nature, 614(7946), 125–135.

Wastyk, H. C., Fragiadakis, G. K., Perelman, D., Dahan, D., Merrill, B. D., Yu, F. B., … & Sonnenburg, J. L. (2021). Gut-microbiota-targeted diets modulate human immune status. Cell, 184(16), 4137–4153.

Wongsurawat, T., Nakagawa, M., Atiq, O., Coleman, H. N., Jenjaroenpun, P., Allred, J. I., … & Nookaew, I. (2019). An assessment of Oxford Nanopore sequencing for human gut metagenome profiling: A pilot study of head and neck cancer patients. Journal of microbiological methods, 166, 105739.

World Health Organization, 1995. Physical status: the use and interpretation of anthropometry. Report of a WHO Expert Committee. Geneva: World Health Organization.

Yatsunenko, T., Rey, F. E., Manary, M. J., Trehan, I., Dominguez-Bello, M. G., Contreras, M., … & Gordon, J. I. (2012). Human gut microbiome viewed across age and geography. Nature, 486(7402), 222–227.

Zeevi, D., Korem, T., Zmora, N., Israeli, D., Rothschild, D., Weinberger, A., … & Segal, E. (2015). Personalized nutrition by prediction of glycemic responses. Cell, 163(5), 1079–1094.

Zhang, C., Zhang, M., Wang, S., Han, R., Cao, Y., Hua, W., … & Zhao, L. (2010). Interactions between gut microbiota, host genetics and diet relevant to development of metabolic syndromes in mice. The ISME journal, 4(2), 232–241.

Zhang, D., Jian, Y. P., Zhang, Y. N., Li, Y., Gu, L. T., Sun, H. H., … & Xu, Z. X. (2023). Short-chain fatty acids in diseases. Cell Communication and Signaling, 21(1), 212.

Zhernakova, A., Kurilshikov, A., Bonder, M. J., Tigchelaar, E. F., Schirmer, M., Vatanen, T., … & Fu, J. (2016). Population-based metagenomics analysis reveals markers for gut microbiome composition and diversity. Science, 352(6285), 565–569.

