## Supplementary Data for "Determinants of gut microbiome composition and its response to a dietary intervention in a multi-ethnic cohort: a pop-up village study"

**List of supplementary material**

| **Material** | **Description** | **Relevant section** |
| --- | --- | --- |
| **Table S1** | Participant metadata | Section 2.1 |
| **Table S2** | Homogeneity of dispersion (betadisper) for each distance metric | Section 3.2 |
| **Table S3** | Timepoint × BMI interaction across three beta-diversity metrics | Section 3.3 |
| **Table S4** | Top five SIMPER contributors in OW participants | Section 3.3 |
| **Table S5** | Multi-tool consensus tiers for whole-cohort differential abundance | Section 3.4 |
| **Table S6** | BMI-stratified core microbiome dynamics | Section 3.4 |
| **Table S7** | Alpha-diversity linear mixed-effects models by BMI subgroup | Section 3.5 |
| **Table S8** | Functional versus taxonomic beta-diversity PERMANOVAs | Section 3.6 |
| **Table S9** | BMI-stratified KO-level MaAsLin2 results by pathway | Section 3.6 |
| **Table S10** | Functional permutation simulation (1,000 paired-swap iterations) | Section 3.6 |
| **Figure S1** | BMI-stratified beta-diversity response to intervention | Section 3.3 |
| **Figure S2** | Taxa contributing to the OW compositional shift (SIMPER) | Section 3.3 |
| **Figure S3** | Genus-level taxa associated with the intervention | Section 3.4 |
| **Figure S4** | Species-level taxa across three association models | Section 3.4 |
| **Figure S5** | Core microbiome turnover by BMI stratum | Section 3.4 |
| **Figure S6** | KO-level functional differential abundance | Section 3.6 |
| **Figure S7** | Functional permutation null distributions and OW exact-pattern analysis | Section 3.6 |

**Supplementary Table S1. Participant metadata.** Participant-level metadata were collected at both baseline and post-intervention timepoints and comprised the following variables: Sample_ID, Participant, Timepoint, date of birth, age, sex, height (cm), body weight (kg), BMI (kg/m², computed as weight / height²), geographic region (International, North India, South India), country of origin (India, International), and habitual diet type (Vegetarian, Non-Vegetarian). The paired analytical cohort comprised 43 participants (86 samples). The cohort was predominantly male (36 male, 7 female), with ages ranging from 20 to 73 years (median 30 years). Baseline BMI ranged from 17.5 to 32.9 kg/m² (mean 24.7 ± 3.4), spanning Non-Overweight to Overweight categories, with 18 participants classified as overweight (BMI ≥ 25 kg/m²) and 25 as non-overweight at baseline. Geographic representation comprised 18 participants from North India, 15 from South India, and 10 international participants. Habitual diet type was approximately balanced, with 25 non-vegetarian and 18 vegetarian participants. All demographic and anthropometric variables were self-reported at enrolment.

**Table S1:** Zugrama_Metadata_Table_S1_csv (a separate csv file is provided)

**Supplementary Table S2. Homogeneity of dispersion (betadisper) for each distance metric.** Multivariate dispersion was compared between baseline (B) and post-intervention (P) samples for each distance. No metric showed a significant difference, indicating that the timepoint effect reported in Table 2 reflects a shift in community centroid rather than increased inter-individual dispersion.

| **Metric** | **F** | **P** | **Disp. B** | **Disp. P** |
| --- | --- | --- | --- | --- |
| **Jaccard** | 1.705 | 0.182 | 0.312 | 0.320 |
| **Unweighted UniFrac** | 1.732 | 0.189 | 0.291 | 0.299 |
| **Bray-Curtis** | 0.909 | 0.367 | 0.544 | 0.535 |
| **Aitchison** | 0.395 | 0.560 | 35.57 | 36.32 |
| **Weighted UniFrac** | 0.059 | 0.806 | 0.106 | 0.105 |

**Supplementary Table S3. Timepoint × BMI interaction across three beta-diversity metrics.** To confirm that the Timepoint × BMI interaction was not an artifact of a single distance metric, we repeated the paired PERMANOVA across three complementary metrics. The interaction was significant for Bray-Curtis and Aitchison distances and directionally consistent, though not significant, for weighted UniFrac, indicating the effect is robust to metric choice.

| **Metric** | **Df** | **SumOfSqs** | **R²** | **F** | **p** |
| --- | --- | --- | --- | --- | --- |
| Bray-Curtis | 1 | 0.296 | 1.16% | 0.998 | 0.021 |
| Weighted UniFrac | 1 | 0.010 | 1.01% | 0.873 | 0.095 |
| Aitchison | 1 | 1,083 | 0.94% | 0.807 | 0.028 |

### **Supplementary Table S4. Top five SIMPER contributors to within-subject Bray-Curtis dissimilarity in OW participants.** SIMPER decomposition on the 34 OW samples attributed 31.4% of total within-subject Bray-Curtis dissimilarity to the top 10 taxa and 45.5% to the top 20. The leading contributors were high-abundance commensals with variable per-participant directionality; *Megamonas funiformis* was the only taxon convergent across the SIMPER ranking, the OW MaAsLin2 model and per-participant directionality.

| **Taxon** | **Avg %** | **Cum %** | **SIMPER p** | **MaAsLin2 q** | **Direction** |
| --- | --- | --- | --- | --- | --- |
| *Megamonas funiformis* | 5.91 | 7.6 | 0.047 | 0.187 | ↓ |
| *Prevotella copri* | 3.01 | 11.5 | 0.602 | 0.829 | ↑ |
| *Coprococcus eutactus* | 3.00 | 15.4 | 0.918 | 0.941 | ↑ |
| *Dialister massiliensis* | 2.20 | 18.3 | 0.060 | 0.896 | ↑ |
| *Faecalibacterium prausnitzii* | 1.95 | 20.8 | 0.037 | 0.581 | ↑ |

### **Supplementary Table S5. Multi-tool consensus tiers for whole-cohort differential abundance.** Whole-cohort differential-abundance results organised by cross-tool consensus. MaAsLin2, LinDA and ALDEx2 were applied to the same filtered feature table; tier assignment reflects the number of tools that identified each taxon at q < 0.25. LinDA identified 21 taxa at padj < 0.05 and 74 at padj < 0.25; ALDEx2, run with an unpaired Welch test and 128 Dirichlet Monte Carlo instances, identified one taxon at q < 0.25 and none at q < 0.05.

| **Tools** | **Count** | **Examples** |
| --- | --- | --- |
| All 3 tools (q < 0.25 each) | 1 | *S. parasanguinis* |
| MaAsLin2 + LinDA (q < 0.25) | 10 | *F. prausnitzii*, *B. adolescentis, Weissella, Megamonas, Lachnospiraceae* bacterium GAM79, *Oscillibacter*, *Phocaeicola, Parabacteroides distasonis* |
| MaAsLin2 only (q < 0.25) | 23 | remaining exploratory taxa |

### **Supplementary Table S6. BMI-stratified core microbiome dynamics.** Core membership was defined as detection at ≥ 0.1% relative abundance in ≥ 50% of participants per timepoint, using the same detection and prevalence thresholds within each subgroup (baseline-BMI assignment). OW-specific post-intervention core entries included butyrate producers (*Agathobaculum butyriciproducens*, *Roseburia hominis*) and fibre degraders (*Bacteroides uniformis*, *B. caccae*, *Phocaeicola vulgatus*). The *Bifidobacterium* split observed in the stratified differential-abundance analysis was recapitulated at the core level: *B. longum* exited the OW core (but remained in the Non-OW core), and *B. adolescentis* exited the Non-OW core (but remained in the OW core).

| **Metric** | **All (n=43)** | **OW (n=18)** | **Non-OW (n=25)** |
| --- | --- | --- | --- |
| Baseline core taxa | 53 | 39 | 56 |
| Post core taxa | 52 | 48 | 50 |
| Stable core taxa | 45 | 28 | 46 |
| Core stability % | 84.9% | 71.8% | 82.1% |
| Taxa entering post | 7 | 20 | 4 |
| Taxa leaving post | 8 | 11 | 10 |
| Net change | −1 | +9 | −6 |
| Jaccard turnover | 0.250 | 0.525 | 0.233 |

### **Supplementary Table S7. Alpha-diversity linear mixed-effects models in the OW and Non-OW subgroups.**

| **Metric** | **OW estimate** | **OW (P)** | **OW (q/FDR)** | **Non-OW (p)** | **Non-OW (q)** | **OW (direction)** |
| --- | --- | --- | --- | --- | --- | --- |
| ***Richness / phylogenetic diversity metrics*** | | | | | | |
| Faith’s PD | -60.10 | 0.023 | 0.071 | 0.84 | 0.897 | ↓ |
| Chao1 | -51.19 | 0.027 | 0.071 | 0.62 | 0.897 | ↓ |
| Observed | -44.25 | 0.037 | 0.071 | 0.9 | 0.897 | ↓ |
| ***Evenness metrics*** | | | | | | |
| Simpson | -0.036 | 0.047 | 0.071 | 0.26 | 0.897 | ↓ |
| Pielou | +0.043 | 0.063 | 0.076 | 0.64 | 0.897 | ↑ |
| Shannon | +0.194 | 0.148 | 0.148 | 0.63 | 0.897 | ↑ |

### **Supplementary Table S8. Functional versus taxonomic beta-diversity PERMANOVAs.**

| **Test** | **Taxonomy (Bray)** | **Function (Bray)** | **Function (filt. TSS)** | **Function (CLR)** |
| --- | --- | --- | --- | --- |
| Timepoint (paired) | p = 0.001 | p = 0.178 | p = 0.378 | p = 0.317 |
| Time × BMI | p = 0.021 | p = 0.231 | p = 0.668 | p = 0.154 |
| Time × Region | P = 0.893 | p = 0.067 | p = 0.961 | p = 0.958 |
| Baseline Region | p = 0.046 | p = 0.051 | p = 0.001 | p = 0.001 |
| Baseline BMI | p = 0.024 | p = 0.523 | p = 0.056 | p = 0.043 |
| Non-OW Timepoint | p = 0.181 | – | – | p = 0.588 |
| OW Timepoint | p = 0.001 | p = 0.133 | – | p = 0.01 |

### **Supplementary Table S9. BMI-stratified KO-level MaAsLin2 results: count of KOs with q < 0.25 within each pathway.** Stratified models showed a marked asymmetry between BMI subgroups: 34 KOs reached q < 0.25 in OW participants versus 4 in Non-OW, an 8.5:1 ratio distributed across all six pathways. The strongest OW signals were in the ammonia pathway (K10672, K14048, K00260, all q < 0.001, all decreased) and the butyrate pathway (K01844, +1.67, q = 0.007), consistent with the taxonomic asymmetry.

| **Pathway** | **OW KOs (q<0.25)** | **Non-OW KOs (q<0.25)** | **Top OW hit** |
| --- | --- | --- | --- |
| **Butyrate** | 9 | 0 | K01844 +1.67 (q = 0.007) |
| **Acetate** | 7 | 0 | K01067 +2.00 (q = 0.031) |
| **Ammonia** | 6 | 0 | K10672 − 0.89 (q < 0.001) |
| **Propionate** | 6 | 1 | K11381 +1.15 (q = 0.079) |
| **Methane** | 5 | 0 | K03389 +1.31 (q = 0.213) |
| **Hydrogen sulfide** | 1 | 3 | K20021 +0.48 (q = 0.239) |
| **Total** | 34 | 4 | 8.5:1 |

### **Supplementary Table S10. Functional permutation simulation (1,000 paired-swap iterations).** A paired-swap permutation simulation tested whether the observed pathway-level directional pattern could arise by chance. In each iteration, baseline and post labels were independently and randomly swapped per participant, and the number of pathways with a positive median log₂ fold-change was tabulated. These results frame the functional directional trends as suggestive rather than definitive.

| **Group** | **Observed pattern** | **Matches** | **Empirical p** |
| --- | --- | --- | --- |
| **Whole cohort (n = 43)** | 6/6 positive | 107/1,000 | 0.107 |
| **Non-OW (n =25)** | 6/6 positive | 154/1,000 | 0.154 |
| **OW (n = 18)** | ≥ 5/6 positive | 256/1,000 | 0.256 |
| **OW (n = 18)** | 5 positive + hydrogen sulfide negative | 61/1,000 | 0.061 |

**Supplementary Figures:**

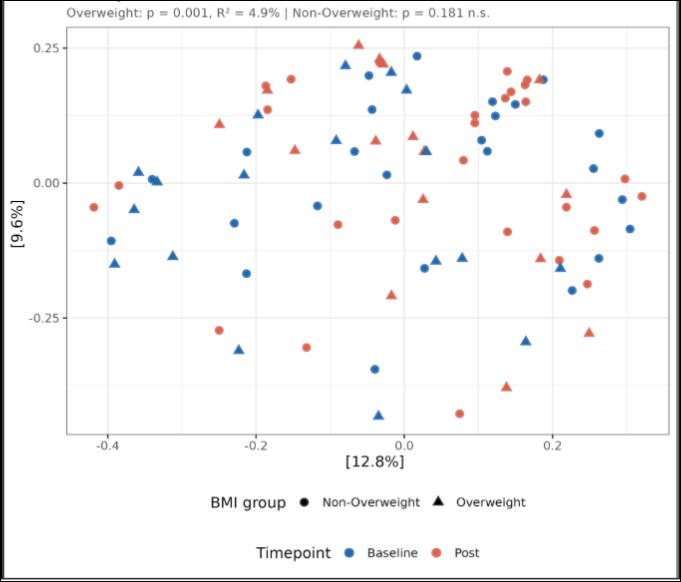

**Fig. S1. BMI-stratified beta-diversity response to intervention.** Single-panel principal coordinates analysis (PCoA) of all 86 paired samples on Bray-Curtis dissimilarity. Marker shape denotes BMI stratum (circles = Non-OW; triangles = OW) and color denotes timepoint (blue = baseline; red = post-intervention).

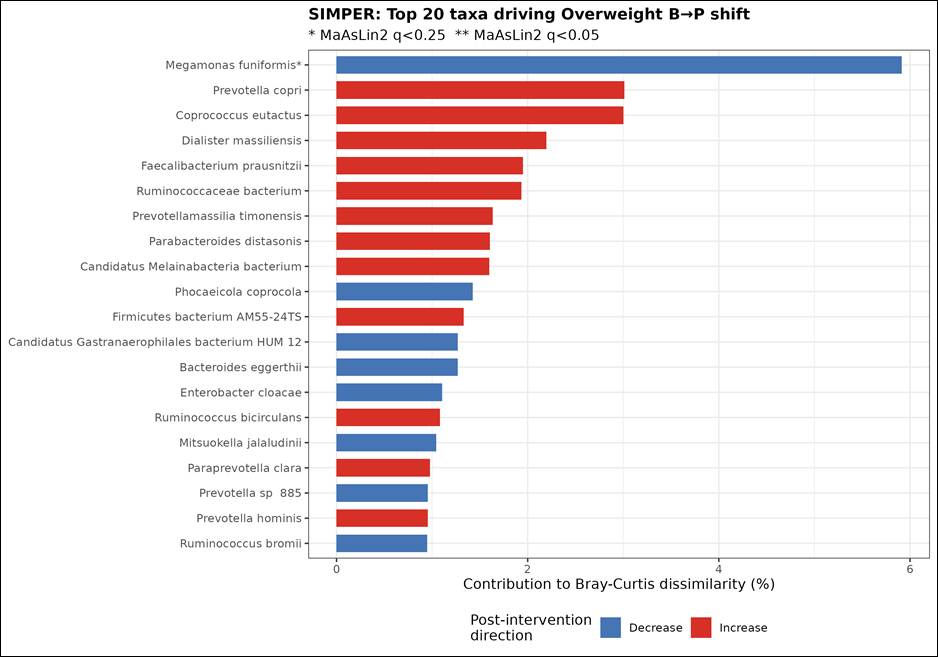
**Fig. S2.** **Taxa contributing to the overweight (OW) compositional shift (SIMPER).** Top 20 taxa ranked by their average percentage contribution to overall Bray–Curtis dissimilarity between baseline and post-intervention OW samples (SIMPER, 999 permutations); bars are colored by direction of change (increased vs decreased post-intervention) and asterisks mark taxa also significant in the MaAsLin2 model (q < 0.25).

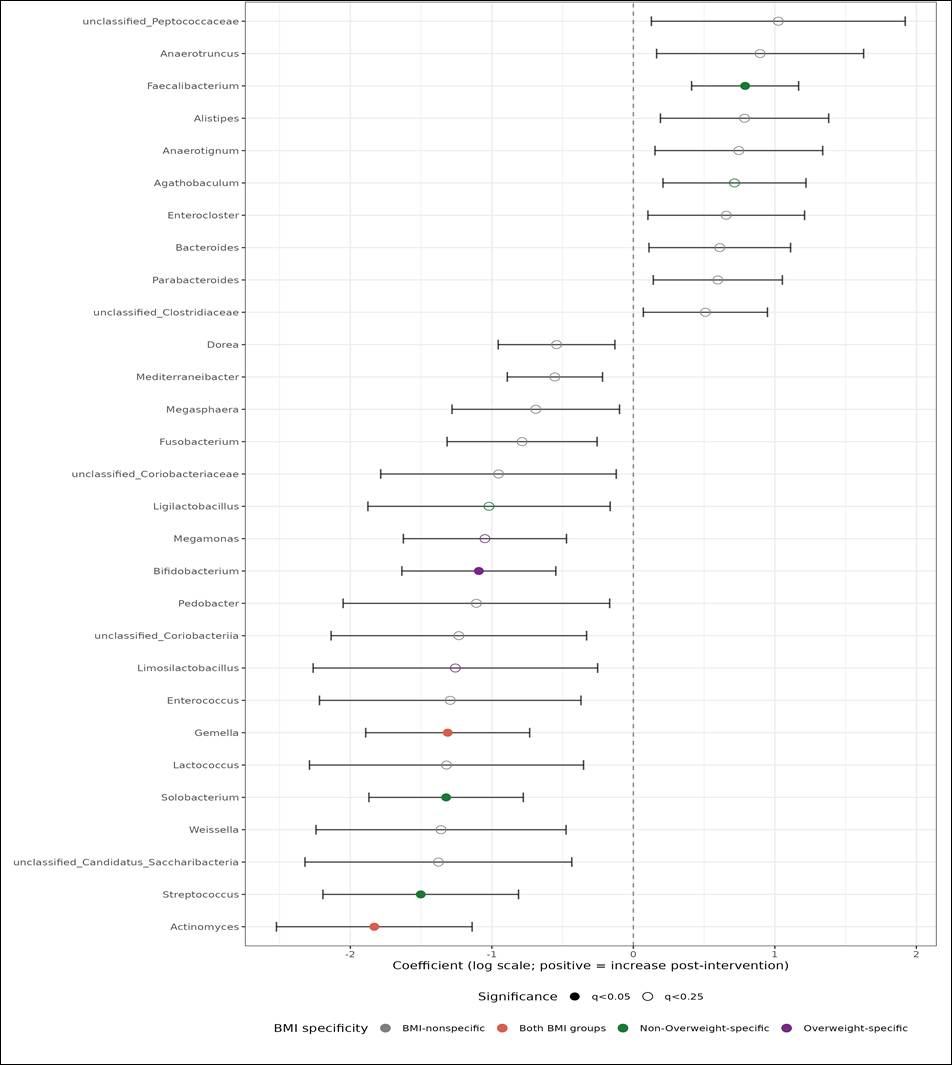
**Fig. S3.** **Genus-level taxa associated with the intervention.** Forest plot of genera differentially abundant between baseline and post-intervention (MaAsLin2, TSS + LOG transform, Participant random effect). Points show model coefficients with 95% confidence intervals; six genera reach q<0.05, led by Actinomyces (q=0.004), Solobacterium (q=0.008) and Gemella (q=0.010). Colour denotes the BMI response pattern of each genus (BMI-nonspecific, both BMI groups, OW-specific, Non-OW specific).

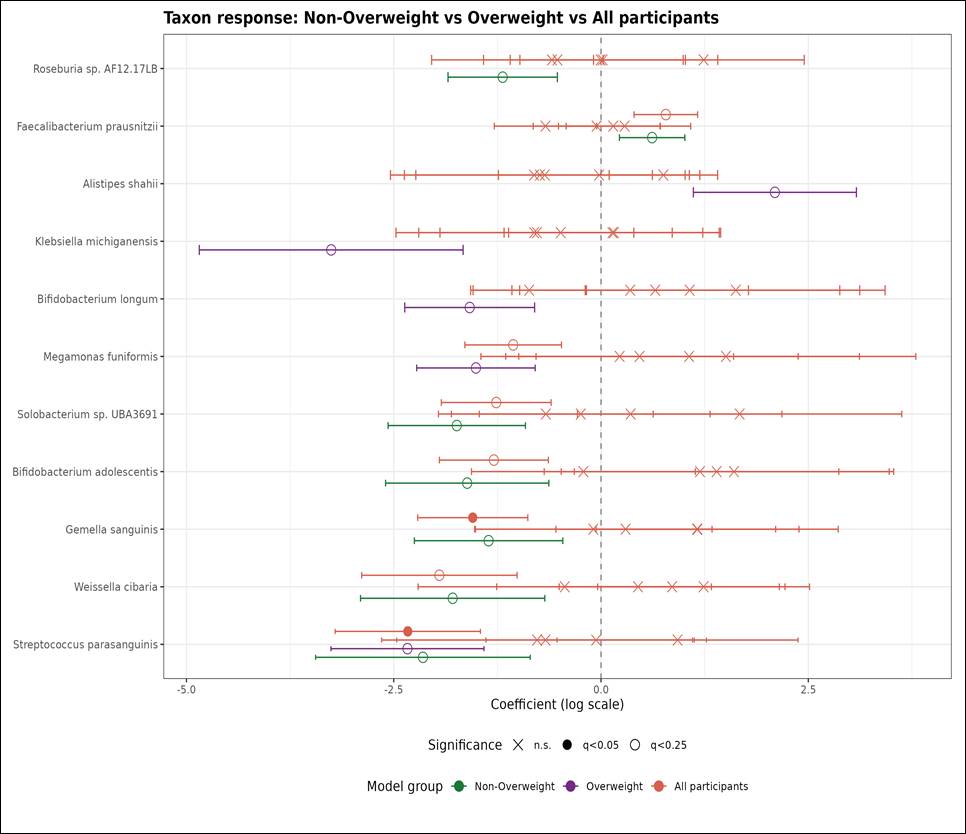
**Fig. S4. Species-level taxa across three association models.** Forest plot of species from three independent MaAsLin2 models: Non-OW samples (52), OW samples (34), and all samples (86). Displayed taxa are the union of species reaching q<0.25 in either stratified model; points show model coefficients with 95% confidence intervals, grouped by model. *Streptococcus parasanguinis* reaches q<0.05 in the all-samples model and q<0.25 in both stratified models.

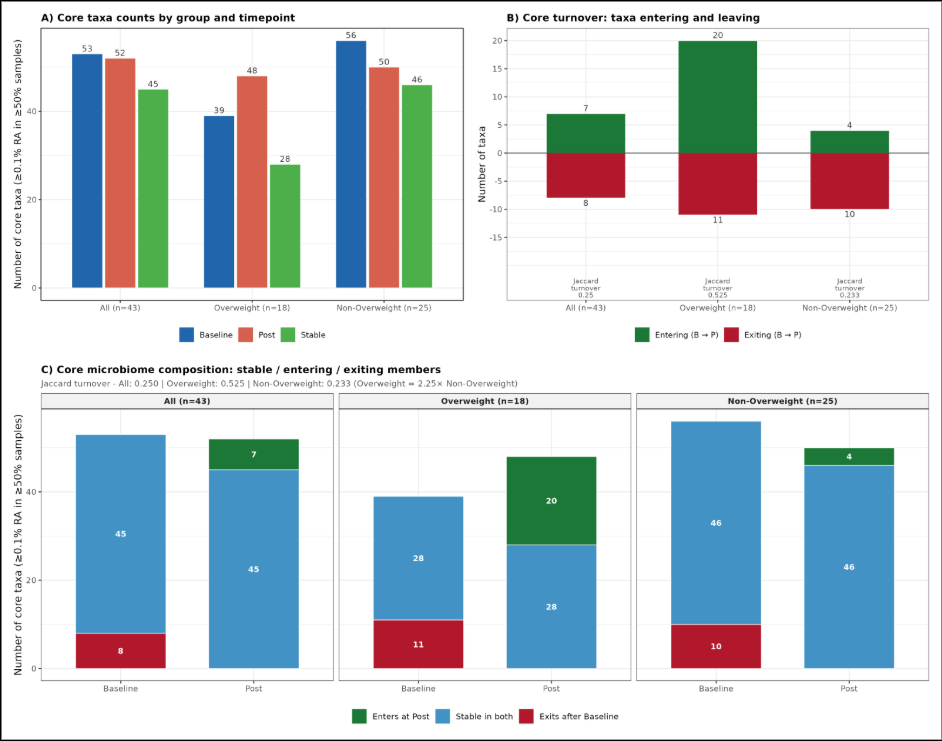
**Fig. S5. Core microbiome turnover by BMI stratum.** **(A)** Core taxa counts at baseline and post-intervention by stratum (OW 39→48 taxa; Non-OW 56→50; All 53→52), and the composition of gained and lost core taxa per stratum. **(B)** Core taxa entering (green) and leaving (red) the core between baseline and post-intervention, with Jaccard turnover annotated, shown for All (n = 43), Overweight (n = 18) and Non-Overweight (n = 25). **(C)** Core composition at each timepoint split into taxa entering at post-intervention (green), stable across both timepoints (blue), or exiting after baseline (red), for the same three groups.

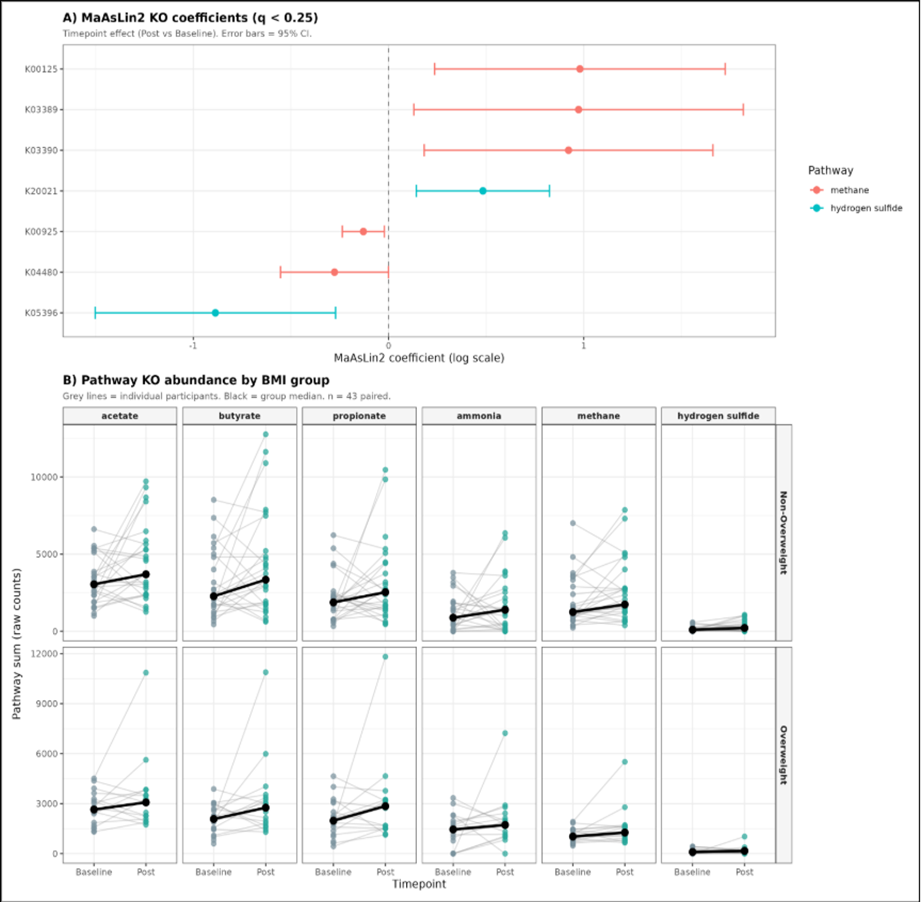

**Fig. S6. KO-level functional differential abundance.** **(A)** MaAsLin2 coefficient forest plot for the seven KEGG orthologs reaching q < 0.25, colored by pathway; points are model coefficients with 95% confidence intervals. **(B)** Paired dot plot of pathway KO relative abundance by BMI subgroup across timepoints (top row, Non-OW; bottom row, OW), each panel showing baseline vs post-intervention.

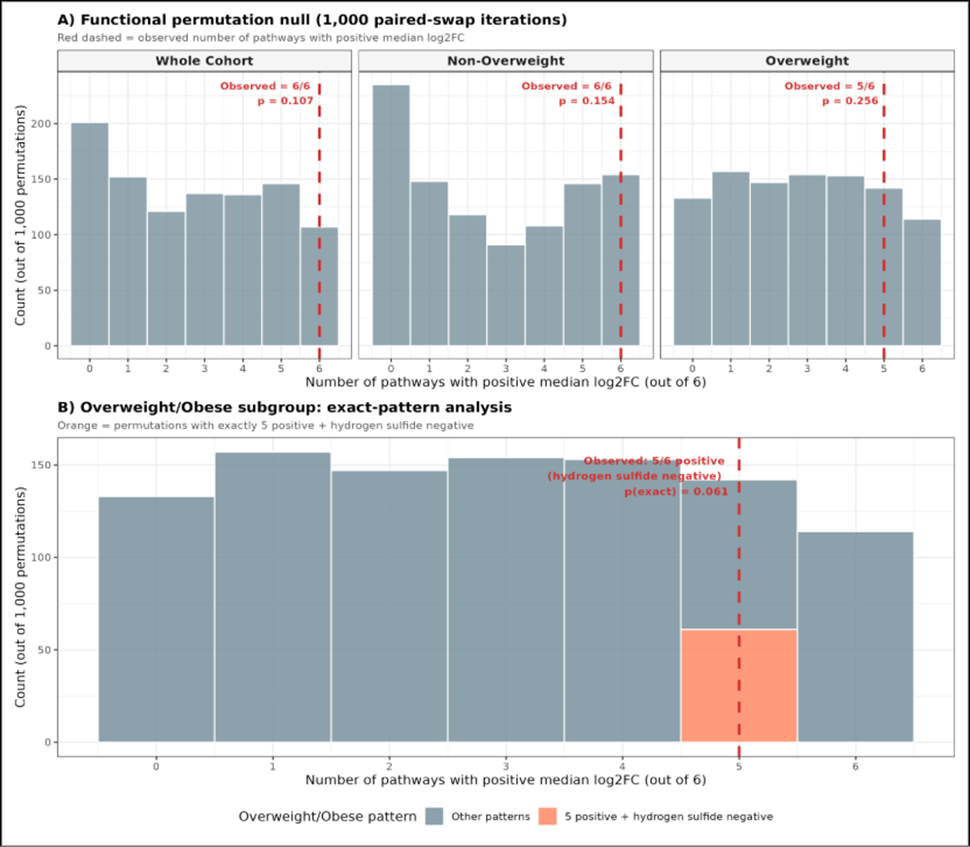

**Fig. S7. Functional permutation null distributions and overweight (OW) exact pattern analysis**. **(A)** Three-facet histograms of the permutation null distribution (1,000 paired-swap iterations) for the number of pathways showing positive directional change, for the whole cohort, Non-OW, and OW strata; the red dashed line marks the observed value in each facet. **(B)** Null distribution for the OW exact-pattern test (5 pathways positive + hydrogen sulfide negative); the orange bar highlights the observed configuration, which occurred in 61/1,000 permutations (p = 0.061).
